# Dimerization of SARS-CoV-2 nucleocapsid protein affects sensitivity of ELISA based diagnostics of COVID-19

**DOI:** 10.1101/2021.05.23.445305

**Authors:** Wajihul Hasan Khan, Nida Khan, Avinash Mishra, Surbhi Gupta, Vikrant Bansode, Deepa Mehta, Rahul Bhambure, Anurag S. Rathore

## Abstract

Diagnostics has played a significant role in effective management of severe acute respiratory syndrome coronavirus 2 (SARS-CoV-2). Nucleocapsid protein (N protein) is the primary antigen of the virus for development of sensitive diagnostic assays. Thus far, limited knowledge exists about the antigenic properties of the N protein. In this paper, we demonstrate the significant impact of dimerization of SARS-CoV-2 nucleocapsid protein on sensitivity of enzyme-linked immunosorbent assay (ELISA) based diagnostics of COVID-19. The expressed purified protein from *E*.*coli* consists of two forms, dimeric and monomeric forms, which have been further characterized by biophysical and immunological means. Indirect ELISA indicated elevated susceptibility of the dimeric form of the nucleocapsid protein for identification of protein-specific monoclonal antibody as compared to the monomeric form of the protein. These findings have also been confirmed with the modelled structure of monomeric and dimeric nucleocapsid protein *via* HHPred software and its solvent accessible surface area, which indicates higher stability and antigenicity of the dimeric type as compared to the monomeric form. It is evident that use of the dimeric form will increase the sensitivity of the current nucleocapsid dependent ELISA for rapid COVID-19 diagnostic. Further, the results indicate that monitoring and maintaining of the monomer-dimer composition is critical for accurate and robust diagnostics.

**Graphical abstract:** 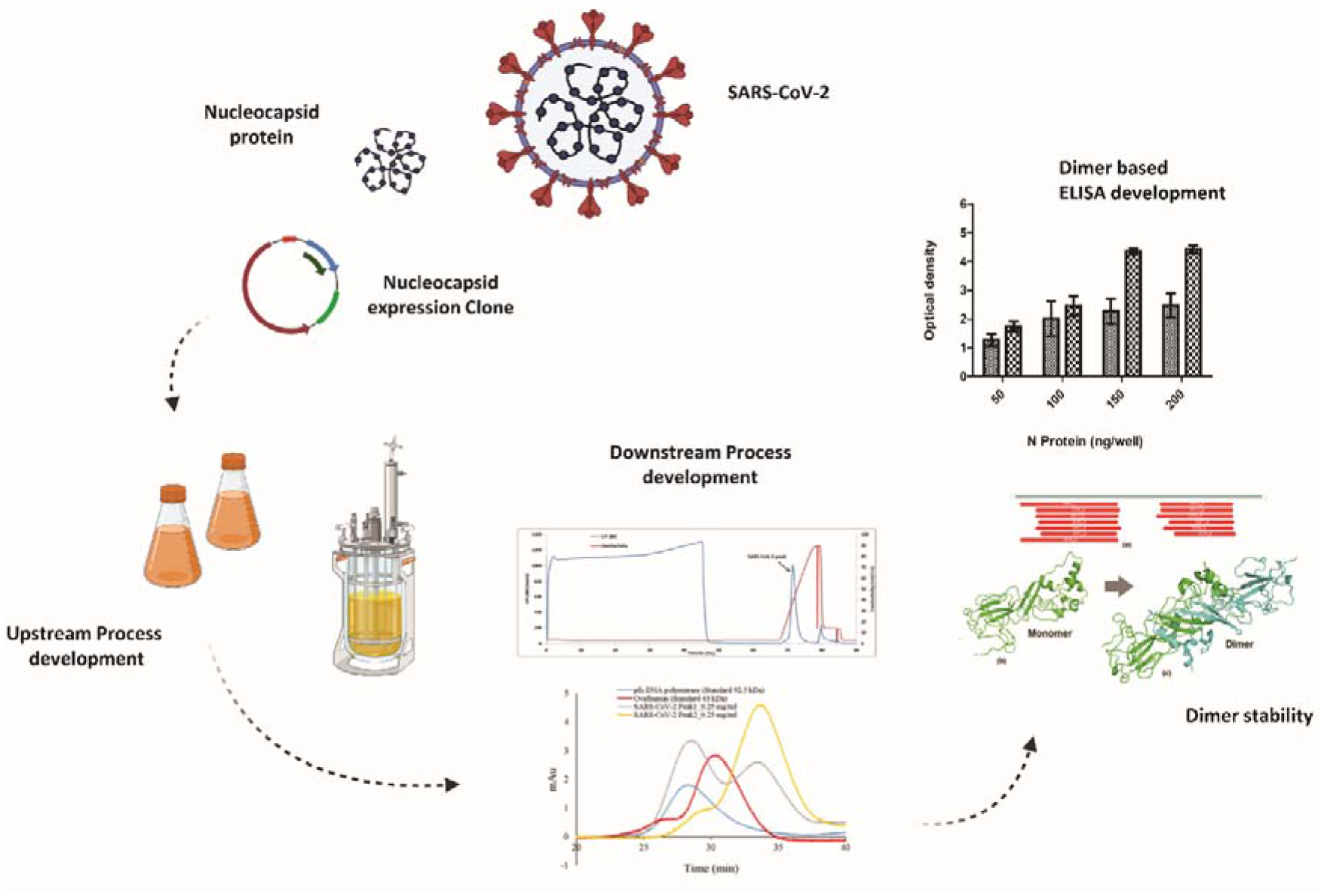

## Introduction

COVID-19 is a widespread global pandemic that has significantly damaged the financial stability and access to treatment for many, especially our most marginalized societies^1-3^. Diagnostics has played a major role in managing the pandemic, with most tests serving as an indicator of transmission at the time when the virus is in the upper respiratory tract^4^. However, detection of pathogen-specific antibodies that develop within days of infection is also a durable biomarker of prior exposure. The antibody-based assay has also been useful in identifying those who have been exposed to the virus^5, 6^.

The SARS-CoV-2 genome is composed of approximately 30,000 nucleotides, which encodes four structural proteins including spike (S) protein, envelope (E) protein, membrane (M) protein, and nucleocapsid (N) protein. SARS-CoV-2 N protein is a ∼45.6 kDa phosphoprotein, comprising of a N-terminal domain (NTD) and a C-terminal domain (CTD), connected by a loosely structured linkage region containing a serine/arginine-rich (SR) domain^7, 8^. The residues from 45 to 181 of the NTD are responsible for the binding of viral RNA to the N protein. SR area linking the NTD and CTD is the site of phosphorylation which is assumed to control N protein performance^9^. Hydrophobic CTD of the N protein contains residues responsible for the homodimerization of the N protein^10-13^. Homodimers of N protein are recorded to self-assemble into higher-order oligomeric complexes, possibly through cooperative interactions of homodimers^14^. Development of higher-order oligomeric complexes requires both dimerization domain and the expanded asymmetric moiety of the CTD^7,15, 16^.

Upon SARS-CoV-2 infection, viral genomic RNA gets associated with the N protein to develop a ribonucleoprotein complex. This complex then packages itself into a helical conformation and combines itself with the M protein of the virion^8^. Despite being present within the viral particle and not very exposed to the surface, SARS-CoV-2 infected patients show elevated and earlier humoral response to the N protein rather than the spike^17^. This is the reason why the N protein is being widely used in vaccine development and serological assays^17-19^. It has been shown for SARS-CoV that the C-terminal region of the N protein is crucial for eliciting antibodies in immunological process^20^. Most diagnostic assays are based on the antigenic proteins, either N or S protein, of the SARS-CoV-2^21-27^. Several formats of ELISA have been developed to detect IgM/IgG antibodies in a patients serum against the SARS□CoV□2 N protein^28, 29^.

Structural study of the full-length coronavirus N protein expressed in *Escherichia coli* is complicated since the recombinant N protein is very susceptible to proteolysis^11^. As a result, minimal information exists on the structure of the SARS-CoV-2 N protein monomer and its assembly into higher-order complexes. In this study, full-length protein of SARS-CoV-2 was successfully expressed in *E. coli* BL21 (DE3) as aggregated inclusion bodies. Two major peaks of the N protein were identified as a monomeric and dimeric conformation *via* size exclusion chromatography coupled with multi-angle static light scattering (MALS), circular dichroism (CD), and fluorescence spectroscopy. Further, the antigenicity of these conformations was compared through a highly sensitive and precise ELISA-based antibody test. The epitope and solvent accessibility of the monomer and dimer forms of the N protein was also predicted using bioinformatics tools to study the structural stability and antigenicity of these conformations. It is evident that use of the dimeric form will increase the sensitivity of the current nucleocapsid dependent ELISA for rapid COVID-19 diagnostic. Further, the results indicate that monitoring and maintaining of the monomer-dimer composition is critical for accurate and robust diagnostics. To the best of our knowledge this is the first indepth investigation into impact of dimerization of SARS-CoV-2 nucleocapsid protein on sensitivity of enzyme-linked immunosorbent assay (ELISA) based diagnostics of COVID-19.

## Results

### Expression and purification of SARS-CoV-2 N protein

Full-length N protein gene construct was transformed into *E. coli* BL21 (DE3) cells. Robust expression of the full-length N protein was observed in the 10 % SDS-PAGE (Figure 1a).

**Figure 1:**
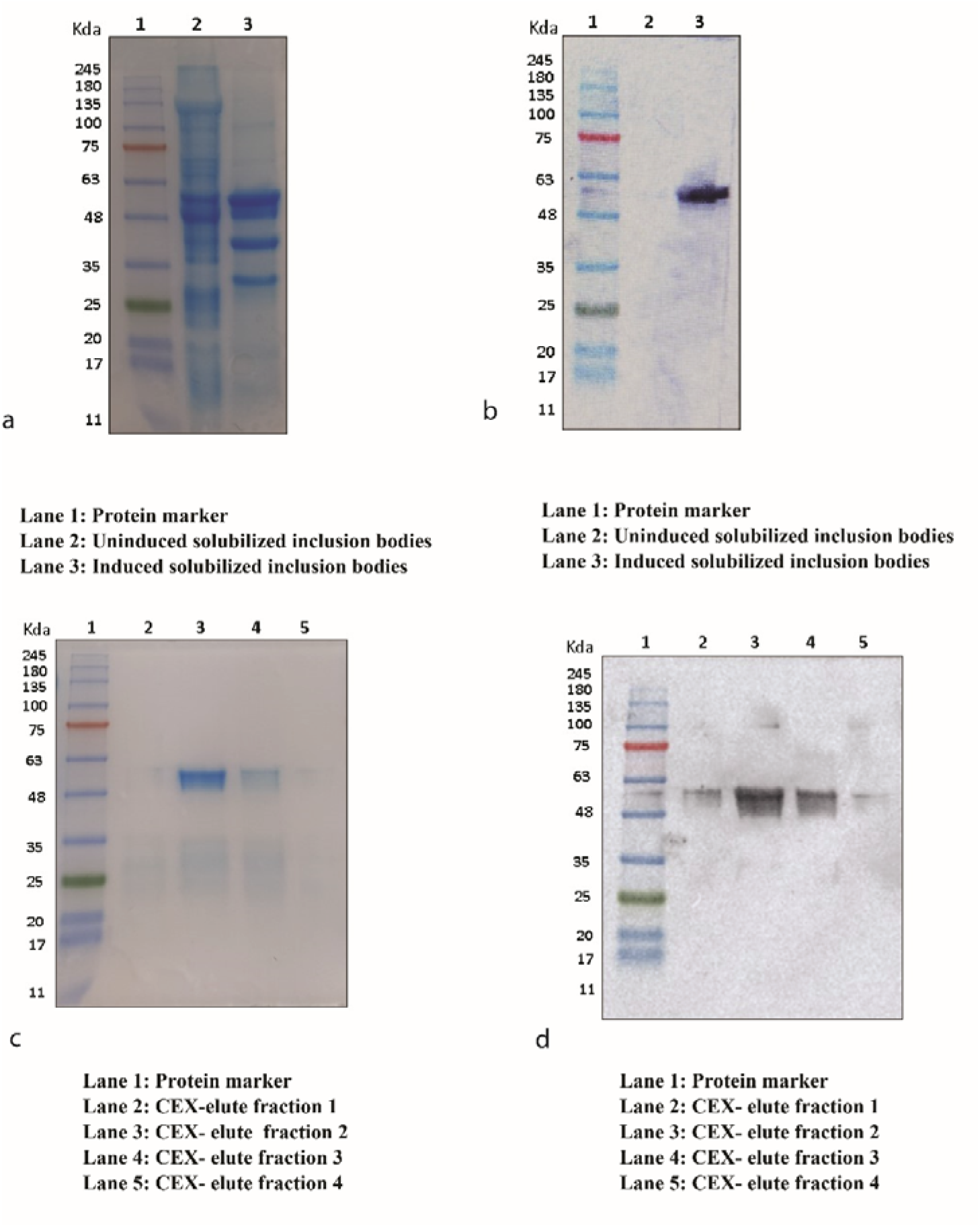
Nucleocapsid protein expression and purification. SDS-PAGE (10%) of expressed N protein **(a)**. Immunoblotting of N protein **(b)**. Coomassie staining of purified N protein fractions **(c)**. Immunoblotting of purified N protein fraction using nucleocapsid specific antibody **(d)**.

The protein band with the molecular weight of about 51.38 kDa represents the full-length N protein expressed as IBs. The protein was further confirmed with immunoblotting using protein-specific antibody (figure 1b).

Protein expression was later scaled-up in a bioreactor and a batch fermentation of transformed *E. coli*. BL21 (DE3) was performed with 10 g L^-1^ (v/v) of glycerol as a carbon source. Upon completion of batch, a DO shoot was observed (Figure 2) and feeding of 200 g L^-1^ (v/v) of the glycerol along with 1% (w/v) yeast extract was given to the bioreactor.

**Figure 2:**
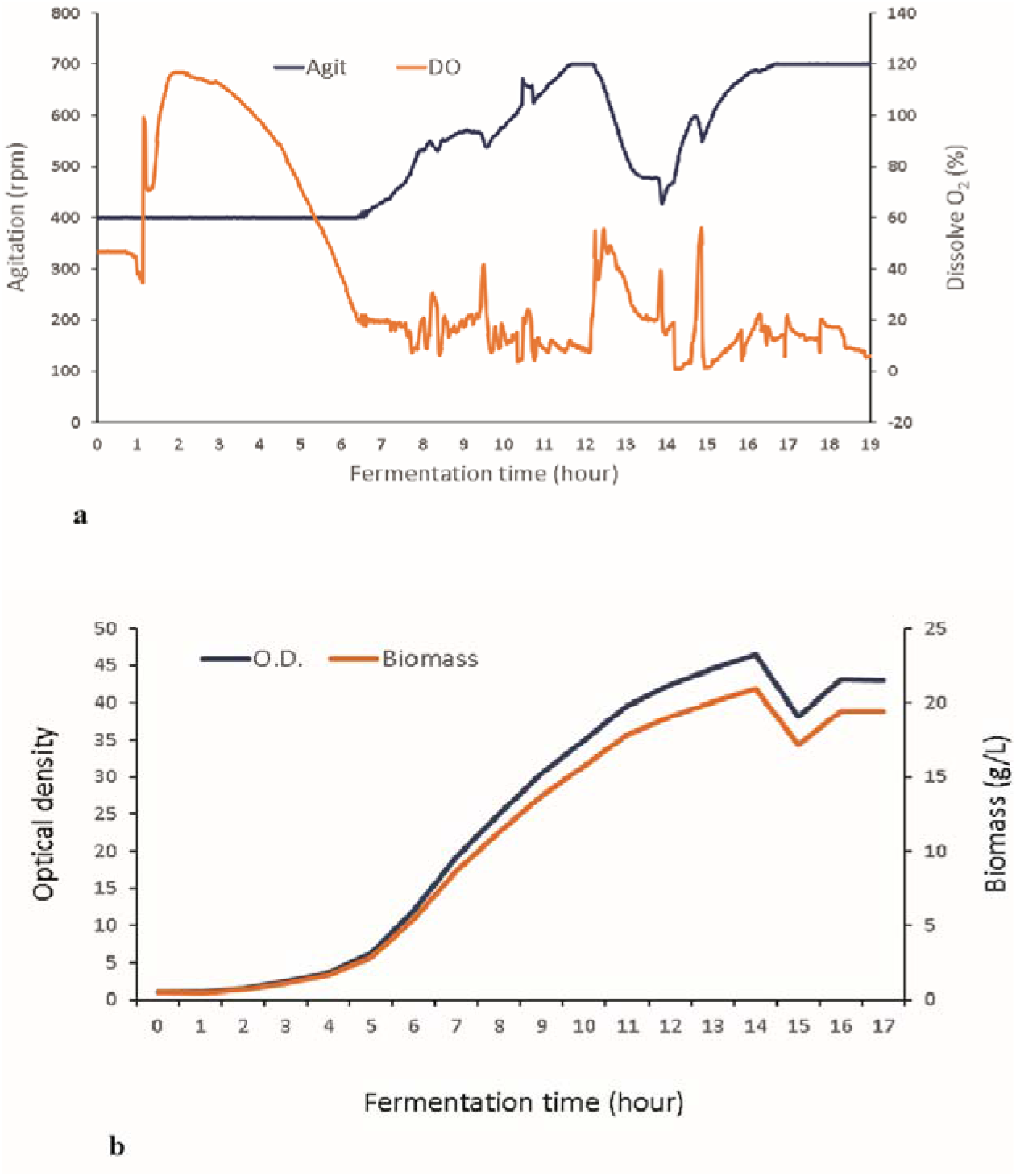
Fermentation profile of nucleocapsid protein production **(a)**. Biomass profile of *E. coli* BL21 (DE3) measured by optical density (OD) **(b)**.

Glycerol feeding resulted in attainment of higher cell density. Protein expression was induced by 1 mM IPTG at an optical density of 35 for 8 h. Biomass of about 20.3 g L^-1^ was generated in the fermentation batch of bioreactor. Due to overexpression of heterologous protein, product was accumulated in the form of IBs within the cytoplasm of the bacterial cell with a yield of about 6.25 g L^-1^.

The inclusion bodies were solubilised, and the protein was captured using SP Sepharose FF resin and purified using CEX chromatography (figure 3). SARS-CoV-2 N protein of more than 95% purity was thus obtained (Figure 1c) and confirmed with immunoblotting (Figure 1d). The protein was conformed with in gel trypsin digestion followed by Liquid Chromatography with mass spectrometry (LC-MS) (figure 4) and SEC-MALS (figure 5) for protein molecular size determination.

**Figure 3:**
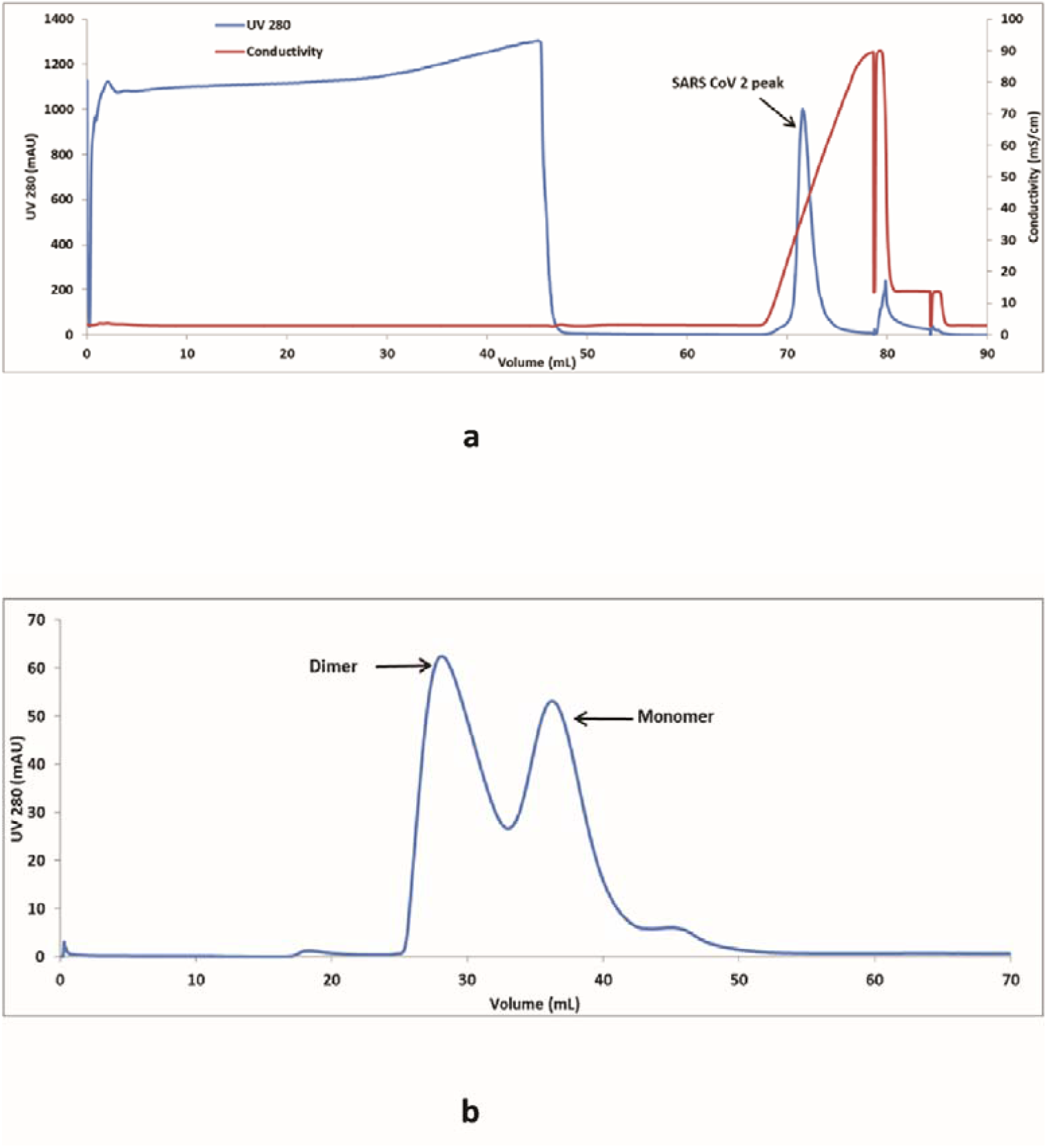
Chromatogram of CEX chromatography **(a)** and preparative SEC **(b)** of Nucleocapsid protein of SARS-CoV-2.

**Figure 4:**
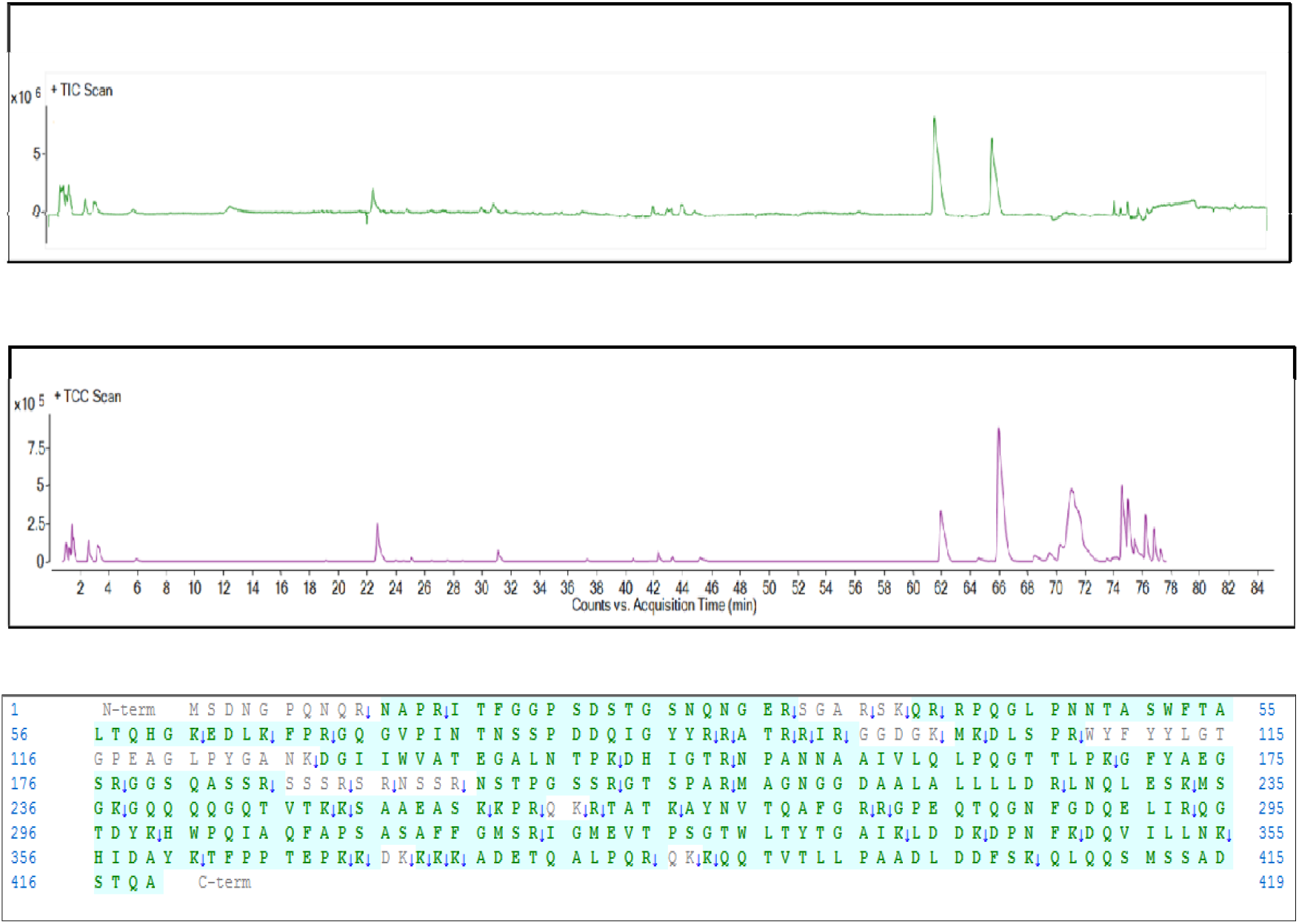
In-gel digestion Mass spectrometry **(a)** total ion chromatogram (TIC), **(b)** total compound chromatogram TCC scan and **(c)** Sequence coverage of trypsin digested SARS-CoV-2 N protein

**Figure 5:**
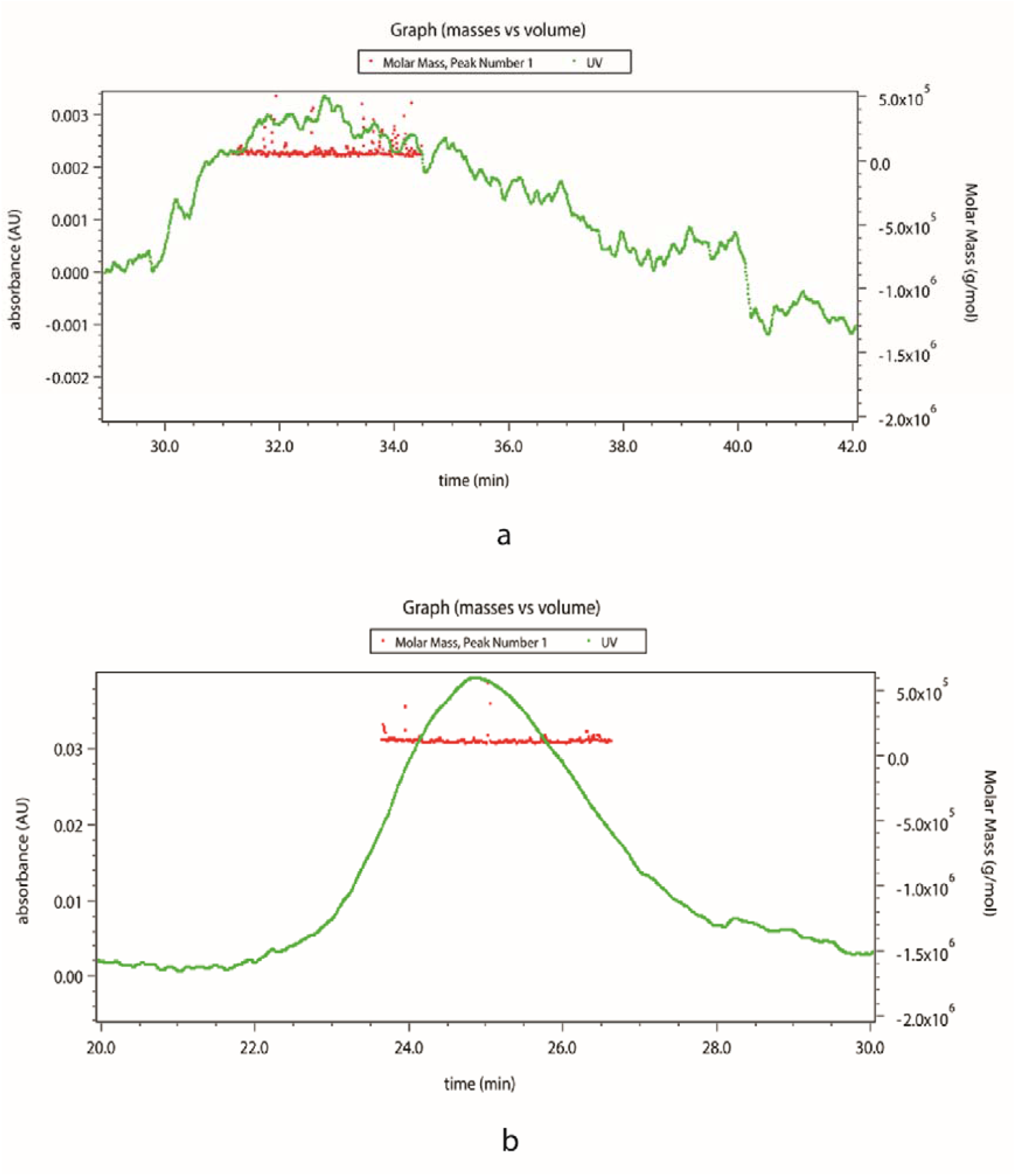
SEC-MALS of SARS-CoV-2 nucleocapsid monomer **(a)** and dimer form of the protein **(b)**.

Further, preparative SEC was performed to obtain fractions containing 5%, 10%, 25%, 55%, 75% dimer (figure 6). Since it is impossible to distinguish the complete dimer from the monomer, fractions with the greatest possible dimer content were used in this analysis. N protein monomer and dimer rich pools were used for structural characterization and determination of ELISA sensitivity.

**Figure 6:**
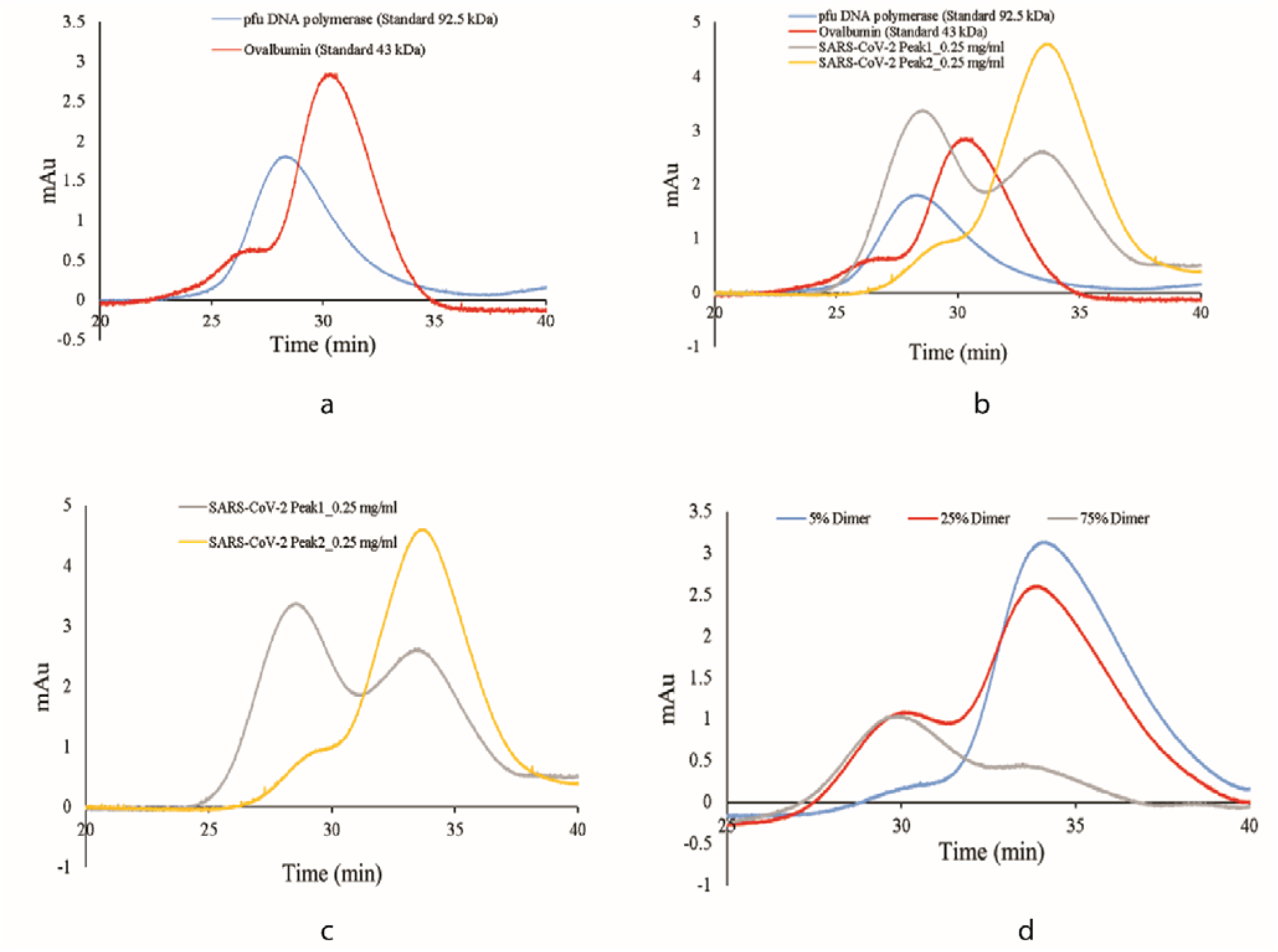
Chromatogram of analytical SEC **(a)** Standard protein of size 43 kDa and 92.5 kDa **(b)** overlay of purified N and standard protein **(c)** Purified N protein alone **(d)** Purified N protein of different fraction of dimer (10%, 25% and 75%)

### Characterization of SARS-CoV-2 N protein

Purified SARS-CoV-2 N protein was analysed by SDS-PAGE, the band was excised and digested with trypsin. The proteolytic digested peptides were analysed using LC-MS. The sequence coverage of the digested protein showed 85.5 % sequence coverage in comparison to in silico digested protein (Figure 4). SEC-MALS and analytical SEC of the N protein showed the monomer and dimer fractions to have molecular masses of 51.38 kDa and 108 kDa respectively (Figure 5 and 6). Further, purified SARS-CoV-2 protein was characterized for secondary structure by CD spectroscopy (Figure 7a). It was observed that SARS-CoV-2 mainly consists of random coils as shown by the negative band at ∼ 200 nm, which is consistent with reports in literature^30^. As is evident from data presented in figure 7b, both the monomer and dimer primarily consist of random coils. In dimer form, there is an increase in ellipticity at 218 nm, as well as a red shift in the negative band from 200 nm to 202 nm, suggesting an improved secondary structure due to oligomerization. Conformational state of SARS-CoV-2 was estimated by fluorescence spectroscopy with tryptophan excitation at 285 nm and emission in the range of 300-500 nm. Fluorescence spectra shows λ_max_ of ∼334 nm (Figure 7b), indicating that the native structure of SARS-CoV-2 protein is similar to that reported for the SARS-CoV N protein^15^. Fluorescence spectra of dimer fraction exhibited a red shift in λ_max_ from 334 nm to 340 nm, indicating exposure of the buried tryptophan and resulting in oligomerization of the protein.

**Figure 7:**
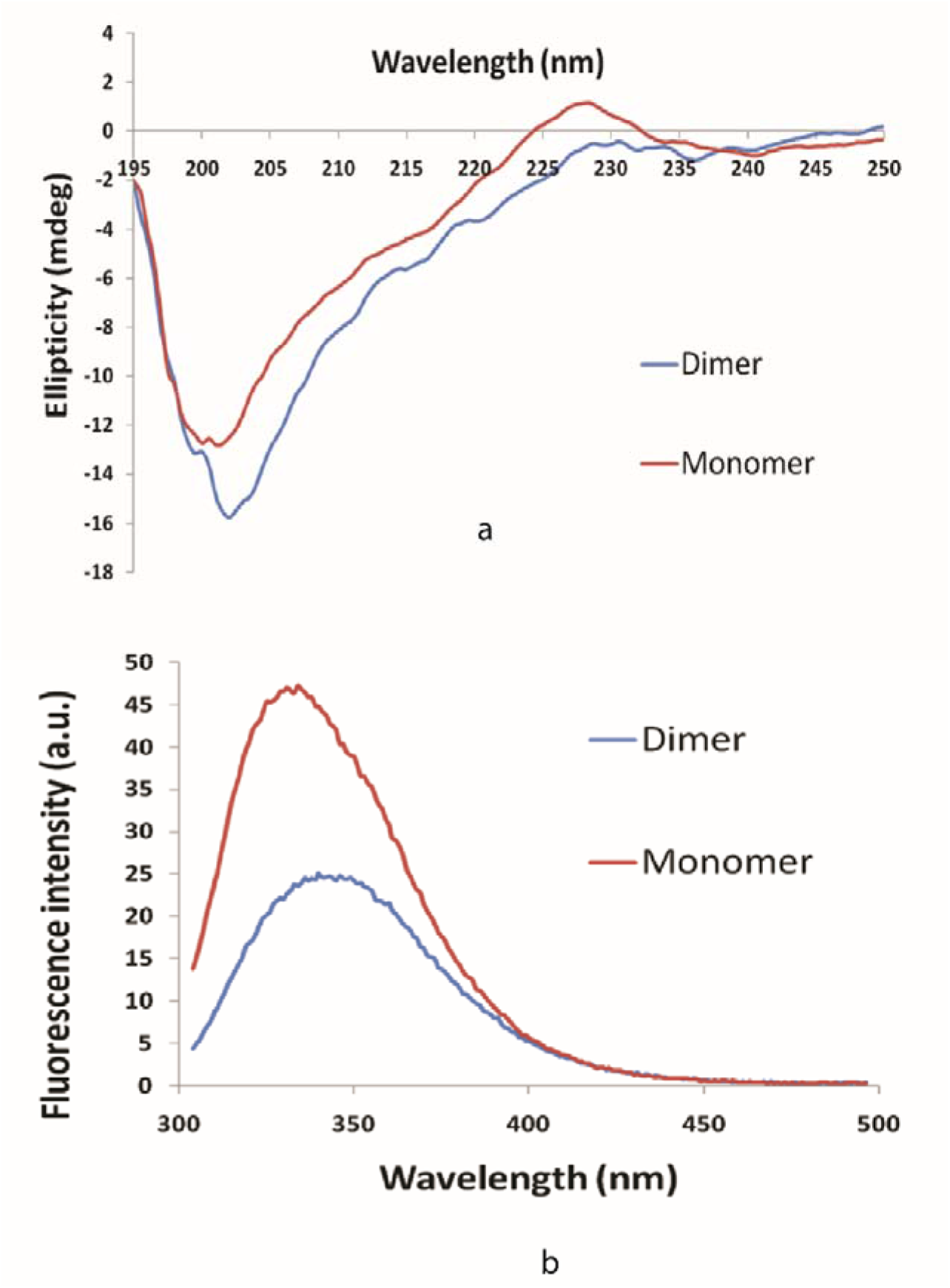
Far-UV Circular Dichroism spectra of N protein of SARS-CoV-2 at 0.25 mg/ml concentration **(a)**. Fluorescence spectra of N protein of SARS-CoV-2 at 0.25 mg/ml concentration **(b)**.

### Modelling of the structure of the monomeric/ dimeric forms

The SARS-CoV-2 N protein sequence was retrieved from the Uniprot database. It comprises of 419 amino acid residues. The current experimental structure contains 30-40% of these residues, rendering it the only structure known for the virus. Sequence alignment showed different potential templates covering the various segments of the protein. Figure 8a shows the coverage of SARS-CoV-2 N protein sequence by different proteins in the sequence alignment. MERS CoV nucleocapsid (PDB: 4UD1) aligned best with the query sequence with 1e-58 E-value and covers 14-164 sequence. Fragment length 244-364 amino acids for SARS-CoV-2 N protein was experimentally solved and used as a second template (PDB: 6WZO). Selected templates (4UD1 and 6WZO) were used in modeller to build the model. This modelled the structure of SARS-CoV-2 N protein in its monomeric form as shown in Figure 8b. Later, this monomeric form was superimposed with the partial experimental structure of the dimerization domain of SARS-CoV-2 N protein (6WZO). Transposing the two units of modelled monomers to the dimerization domain resulted in formation of the dimeric form of the protein as shown in figure 8c.

**Figure 8:**
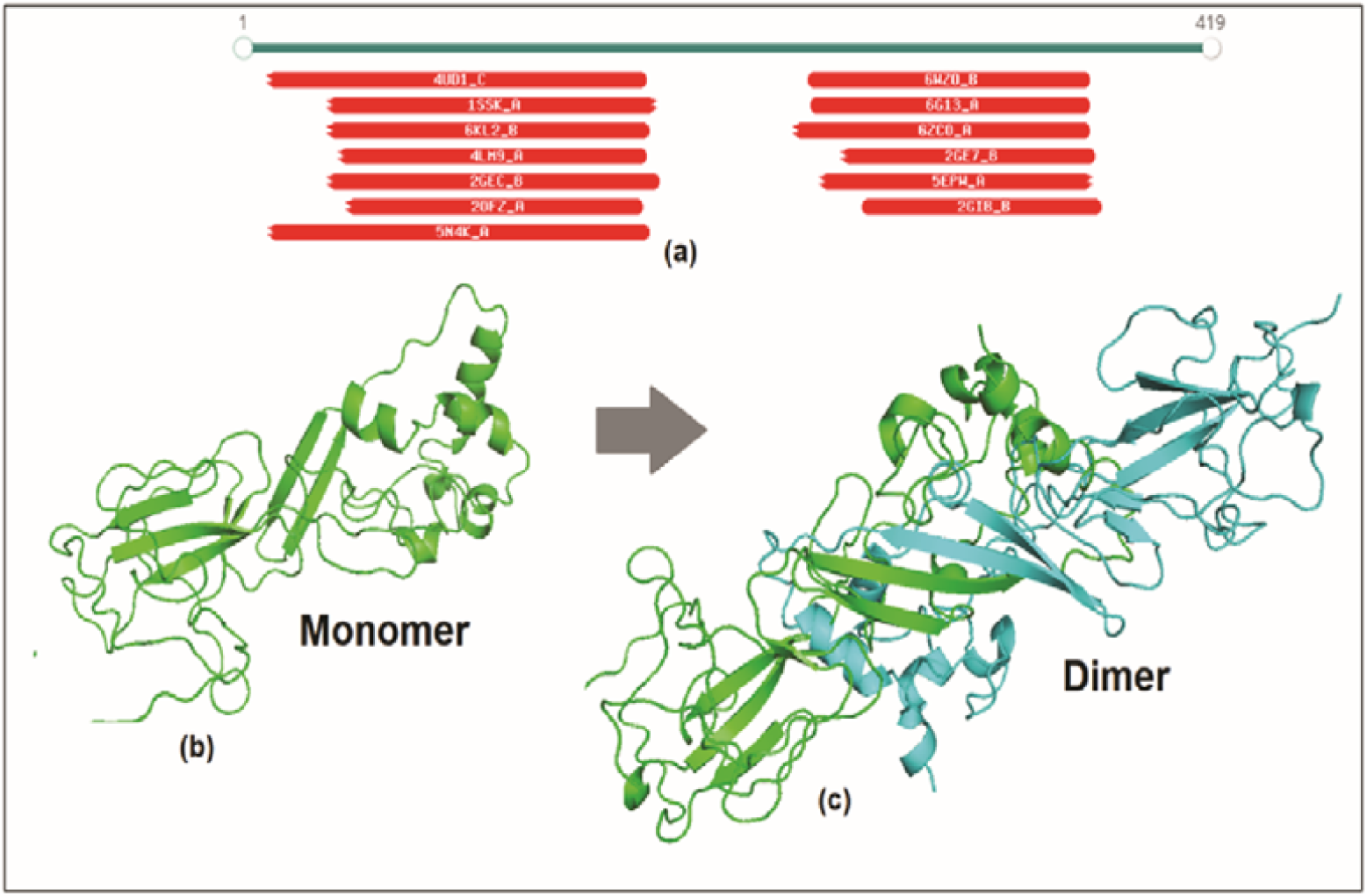
Structure modelling of SARS-CoV-2 N protein **(a)** sequence alignment of SARS-CoV-2 N protein with known structures **(b)** monomer modelled structure built on 4UD1 and 6WZO templates **(c)** dimeric form of protein built using 6WZO dimerization domain.

### Calculation of the solvent accessible surface area (SASA)

SASA was calculated for each residue in monomer and dimer form. This indicates the amount of area for a residue that is exposed to the solvent. Hydrophobic residues do not prefer polar environment and thus bury themselves in the native structure of the protein. Hydrophobic residues that show>50Å2 SASA value in monomer indicate probable surface instability. These residues were marked and their corresponding percentage change in dimeric form was calculated. Figure 9 shows the percentage change of these hydrophobic residues between the monomer and the dimer. Six such residues, namely A183, V182, I283, L271, I269, and I289, changed from completely exposed to buried state in dimeric form. Moreover, A268 buried by 60% while A243 and I194 buried by ≈ 35% compared to their monomeric conformation. This indicates that dimerization of protein helped to bury these hydrophobic residues at the interface that can stabilize the protein in the solution.

**Figure 9:**
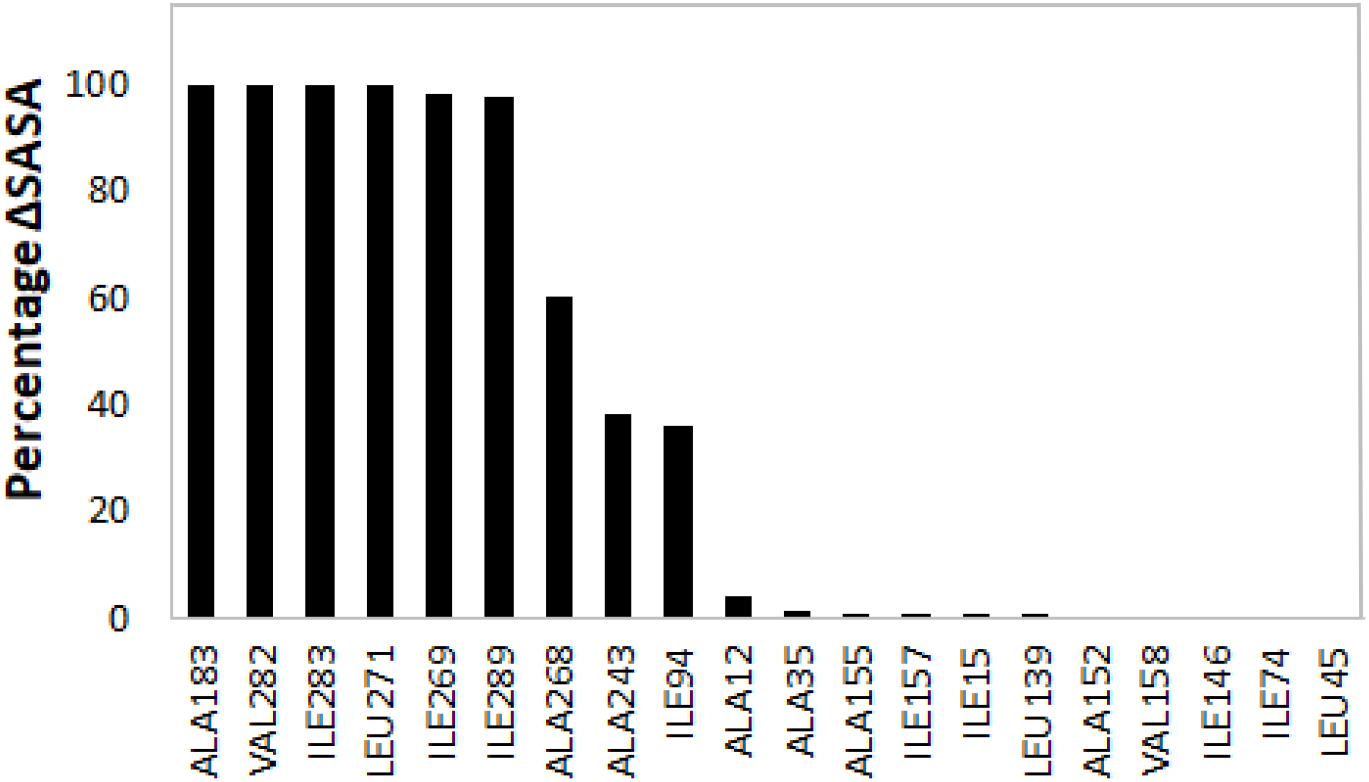
Percentage change in solvent accessible surface area (SASA) for hydrophobic residues between monomer and dimer form of SARS-CoV-2 N protein.

### Prediction of epitopes

The 3D structure of a protein can be used to predict discontinuous epitopes. These epitopes are formed due to specific conformation of protein residues at the surface. To classify these discontinuous epitopes, many methods are used to evaluate monomer and dimer. Ellipro predicted 3 epitopic sites on the monomer surface and 6 epitopes for the dimer form. These 6 predicted epitopes for the dimer SARS-CoV-2 N are the duplicate of its corresponding monomer and hence improve the chance of antibody binding.

Among the predicted discontinuous 3D epitopes, patch 1:”R41, P42, Q43, G44, L45, P46, N47, N48, T49, A50, S51, W52, F53, T54, A55, E62, D63, L64, K65, F66, P67, G69, Q70, G71, V72, P73, I74, N75, T76, N77, S78, S79, P80, D81, D82, Q83, I84, Y112, L113, G114, T115, P122, Y123, G124, A125, V133, A134, T135, E136, G137, A138, L139, N140, T141, P142, K143, D144, H145, I146, G147, T148, R149, N150, P151, A152, N153, N154, A155, A156, I157, V158, L159, Q160, L161, P162, Q163, G164, T165, T166, L167, P168, K169, Y172, A173, E174, G175, Q176, T177, T257, P258, S259, G260” has the maximum score and it is present in duplicate for the dimeric form. A complete list of the epitopes predicted by Ellipro is given in Table 1. Further, a similar analysis was performed with the DiscoTope server, which predicted the probability of each residue to be part of an epitope. It predicted 110 residues at the epitopic site at the DiscoTope threshold score “0”. However, dimer has 238 B-cell epitope residues out of 576 residues. A list of the predicted epitope residues is shown in Supplementary Table S1. Both the servers suggested that dimer has a greater number of structural epitopes and may have more affinity for antibodies.

**Table 1.** Discontinuous epitopes predicted by Ellipro server for the monomeric and dimeric form of SARS-CoV-2 N protein.

| MONOMER (Chain A) |  |  |
| --- | --- | --- |
| Residues | Number of residues | Score |
| A:R194, A:T195, A:A196, A:T197, A:K198, A:A199, A:Y200, A:N201, A:T203, A:Q204, A:G207, A:R208, A:R209, A:G210, A:P211, A:E212, A:Q213, A:T214, A:Q215, A:N217, A:G219, A:D220, A:Q221, A:E222, A:L223, A:I224, A:R225, A:Q226, A:G227, A:T228, A:D229, A:Y230, A:K231, A:H232, A:W233, A:P234, A:Q235, A:I236, A:A237, A:Q238, A:F239, A:A240, A:P241, A:S242, A:S244, A:A245, A:G248, A:M249, A:S250, A:G267, A:A268, A:I269, A:K270, A:L271, A:D272, A:D273, A:K274, A:D275, A:P276, A:N277, A:F278, A:K279, A:D280, A:Q281, A:V282, A:I283, A:L284, A:L285, A:N286, A:K287, A:D290, A:A291, A:Y292, A:K293, A:T294, A:F295, A:P296 | 77 | 0.645 |
| A:F53, A:T54, A:E62, A:D63, A:L64, A:K65, A:F66, A:P67, A:G69, A:Q70, A:G71, A:V72, A:P73, A:I74, A:N75, A:T76, A:N77, A:S78, A:S79, A:P80, A:D81, A:D82, A:Q83, A:A134, A:T135, A:E136, A:G137, A:A138, A:L139, A:N140, A:T141, A:R149, A:N150, A:P151, A:A152, A:N153, A:N154, A:A155, A:A156, A:I157, A:V158, A:L159, A:Q160, A:L161, A:P162, A:Q163, A:G164, A:T165, A:T166, A:L167, A:P168, A:K169 | 52 | 0.613 |
| A:G96, A:G97, A:D98, A:G99, A:K100, A:M101, A:K193 | 7 | 0.505 |
| DIMER (Chain A and B) |  |  |
| Residues | Number of residues | Score |
| A:R41, A:P42, A:Q43, A:G44, A:L45, A:P46, A:N47, A:N48, A:T49, A:A50, A:S51, A:W52, A:F53, A:T54, A:A55, A:E62, A:D63, A:L64, A:K65, A:F66, A:P67, A:G69, A:Q70, A:G71, A:V72, A:P73, A:I74, A:N75, A:T76, A:N77, A:S78, A:S79, A:P80, A:D81, A:D82, A:Q83, A:I84, A:Y112, A:L113, A:G114, A:T115, A:P122, A:Y123, A:G124, A:A125, A:V133, A:A134, A:T135, A:E136, A:G137, A:A138, A:L139, A:N140, A:T141, A:P142, A:K143, A:D144, A:H145, A:I146, A:G147, A:T148, A:R149, A:N150, A:P151, A:A152, A:N153, A:N154, A:A155, A:A156, A:I157, A:V158, A:L159, A:Q160, A:L161, A:P162, A:Q163, A:G164, A:T165, A:T166, A:L167, A:P168, A:K169, A:Y172, A:A173, A:E174, A:G175, A:Q176, A:T177, A:T257, A:P258, A:S259, A:G260 | 92 | 0.68 |
| B:R41, B:P42, B:Q43, B:G44, B:L45, B:P46, B:N47, B:N48, B:T49, B:A50, B:S51, B:W52, B:F53, B:T54, B:A55, B:E62, B:D63, B:L64, B:K65, B:F66, B:P67, B:G69, B:Q70, B:G71, B:V72, B:P73, B:I74, B:N75, B:T76, B:N77, B:S78, B:S79, B:P80, B:D81, B:D82, B:Q83, B:I84, B:Y112, B:L113, B:G114, B:T115, B:P122, B:Y123, B:G124, B:A125, B:V133, B:A134, B:T135, B:E136, B:G137, B:A138, B:L139, B:N140, B:T141, B:P142, B:K143, B:D144, B:H145, B:I146, B:G147, B:T148, B:R149, B:N150, B:P151, B:A152, B:N153, B:N154, B:A155, B:A156, B:I157, B:V158, B:L159, B:Q160, B:L161, B:P162, B:Q163, B:G164, B:T165, B:T166, B:L167, B:P168, B:K169, B:Y172, B:A173, B:E174, B:G175, B:Q176, B:T177, B:T257, B:P258, B:S259, B:G260 | 92 | 0.678 |
| A:R194, A:T195, A:A196, A:T197, A:Y200, A:N201, A:T203, A:Q204, A:G207, A:R208, A:R209, A:G210, A:P211, A:E212, A:Q213, A:T214, A:Q215, A:G216, A:N217, A:D220, A:Q221, A:R225, A:Q226, A:D229, A:Y230, A:D290, A:Y292, A:K293, A:T294, A:F295, A:P296 | 31 | 0.529 |
| B:R194, B:T195, B:A196, B:T197, B:Y200, B:N201, B:T203, B:Q204, B:G207, B:R208, B:R209, B:G210, B:P211, B:E212, B:Q213, B:T214, B:Q215, B:G216, B:N217, B:D220, B:Q221, B:R225, B:Q226, B:D229, B:Y230, B:D290, B:Y292, B:K293, B:T294, B:F295, B:P296 | 31 | 0.528 |
| A:D272, A:D273, A:K274 | 3 | 0.508 |
| B:D272, B:D273, B:K274 | 3 | 0.508 |

### Enzyme-linked immunosorbent assay

Antigenic response of the monomeric and dimeric form of the N protein was evaluated through indirect ELISA using monoclonal antibody targeted against nucleocapsid of SARS-CoV-2. In order to improve the assay performance, a systemic perusal of each step of the ELISA was performed. Variables such as antigen-coating concentration and primary antibody dilution were optimized. Four concentrations of N protein monomer and dimer fractions (50-200 ng well^-1^) were coated on the 96-well polystyrene plate. As illustrated in Figure 10b, at a protein concentration of 150 ng well^-1^, dimeric fraction of the N protein was clearly superior for anti-N IgG detection. Following antigen-coating, an ideal primary antibody dilution was tested. Six-two-fold dilutions of primary antibody were generated for the detection of coated N-protein monomer and dimer. Initially, antigen detection was increased with the increase in primary antibody dilution. The detectability saturated at a dilution of 1:3000 dilution and upon further increase in the dilution of the primary antibody, the detection worsened as shown in Figure 10a. Owing to the superior antigenicity of the dimeric form of the N protein, impact of increasing percentage of dimer in the solution on its immunogenicity was further examined. Samples containing different percentages of N protein dimers (10-75%) were coated on the polystyrene plate and were detected by primary antibody (anti-N IgG, 1:3000). It can be inferred from Figure. 10b and c that as the percentage of the dimers increased in the solution, the sensitivity of the assay increased as well (p value <0.05).

**Figure 10:**
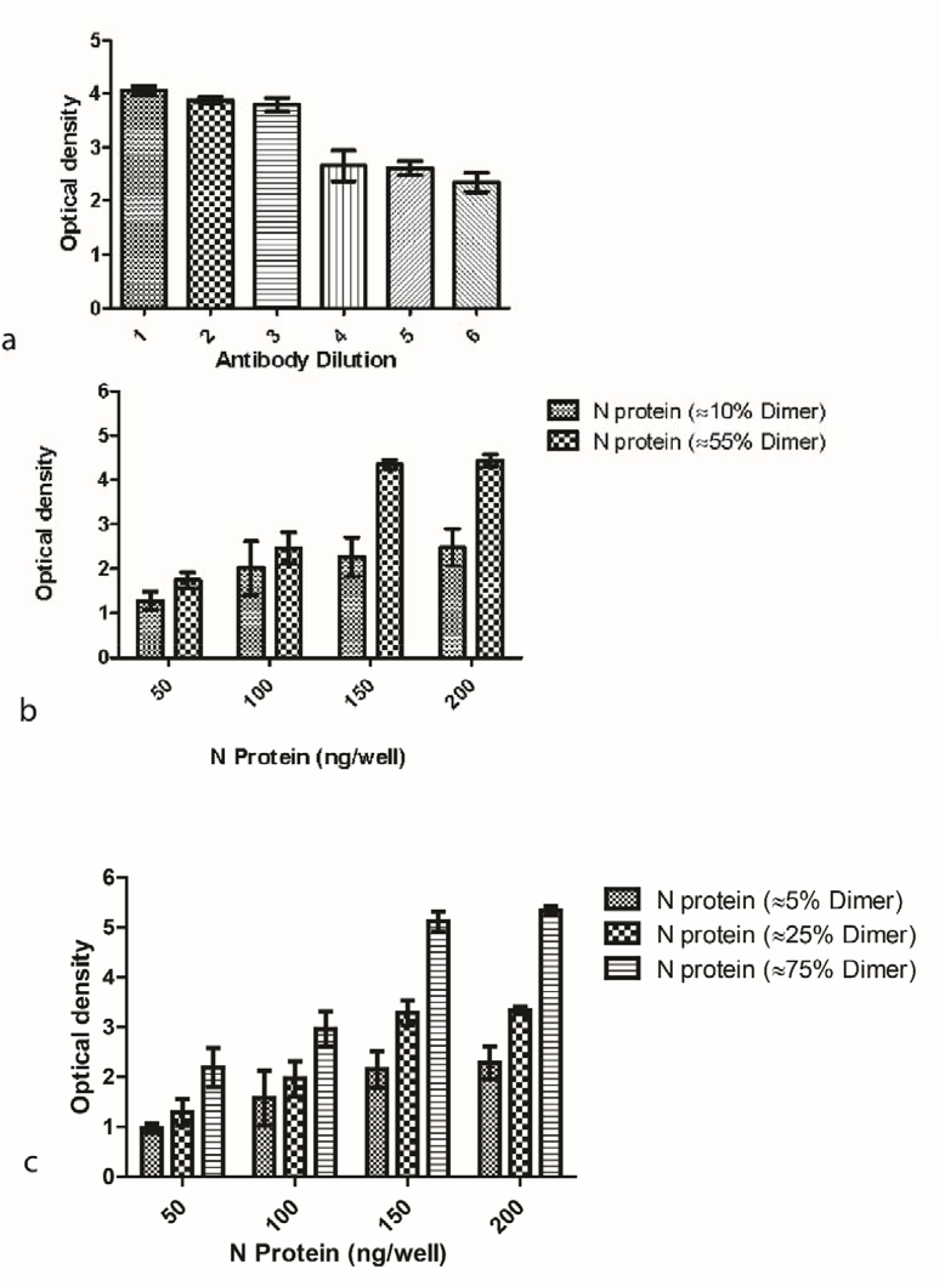
Indirect ELISA based on N protein. **(a)** Standardization of antibody dilution **(b)** ELISA based on increasing amount of N protein dimer of fraction 10% and 55% **(c)** ELISA on increasing amount of N protein dimer fraction of 5%, 25% and 75%.

## Discussion

The SARS-CoV-2 nucleocapsid protein antibody is more sensitive than the spike protein antibody for ELISA based identification of early infections^31^. N protein is a highly immunogenic and generously expressed protein during infection of SARS-CoV-2. High levels of anti-N protein antibodies have been detected in sera in patients with prior infection of SARS-CoV-2. SARS-CoV-2 N protein is a highly basic protein with a pI of 10.0. The nucleocapsid is a multifunctional protein that interacts with RNA and other membrane proteins during virus assembly. Researchers have reported that homodimers of the full-length N protein are the fundamental unit of the ribonucleoprotein complex^32^.

In the present study, we purified monomeric and dimeric form of the nucleocapsid protein. The dimer and monomer ratio did not change in the concentration range under consideration. Samples rich in monomeric or dimeric forms were used to investigate the antigenic sensitivity to the SARS-CoV-2 nucleocapsid. The highest-grade dimer fraction of the N protein demonstrated high sensitivity and a wider dynamic range for antibody detection. Later, we investigated the phenomenon of high sensitivity of dimer using computational approach. Structures of N protein were modelled in their monomeric and dimeric forms. Solvent accessibility of hydrophobic residues was found to be lower in the dimeric form, thereby indicating better surface stability for dimer in polar solution. Additionally, 3D epitopes were predicted to find the potential of binding of the monomeric and the dimeric species with the antibody. Dimer species exhibited double the number of epitopes compared to the monomer, thereby enhancing the chances to interact with the antibody, an observation supported by experimental data.

This is the first of its kind study, elucidating the impact of dimerization of SARS-CoV-2 nucleocapsid protein on sensitivity of enzyme-linked immunosorbent assay (ELISA) based diagnostics of COVID-19. The optical density calculation in the ELISA assay improved when a high proportion of the dimeric fraction was used as antigen. Thus, further modification of existing assays for the detection of SARS-CoV-2 antibodies and use of a high proportion of full-length nucleocapsid fragment dimer can further enhance the sensitivity of the existing rapid kit and ELISA assay.

## Methods

### Construct and expression of SARS-CoV-2 N protein

*Escherichia coli* BL21 (DE3) purchased from Novagen – Merck Life Science Private Limited, India (Cat. No.69450-4), was used in the current study. The expression construct of SARS-CoV-2 N protein was commercially procured through Addgene in a pGBW-m4046785 vector with N protein gene insert of 1253 bp under the control of T7 promoter. The expression construct was transformed in *E*.*coli* BL21 (DE3) strain and bacterial culture was grown in terrific broth at 37.0 °C in the presence of 25 µg mL^-1^ chloramphenicol. When the O.D. at 600 of primary culture reached up to 1.0 ± 0.2, the secondary culture (100.0 ml) was inoculated with 5.0 ml of primary seed culture. Bacterial cultivations were carried out at 37.0 °C. 1.0 mM isopropyl β□D□1□thiogalactopyranoside (IPTG) was used to induce the secondary culture in the midlog phase. Cells were harvested after 12 hours of induction and subjected to primary downstream processing steps to confirm protein expression.

### Production of SARS-CoV-2 N protein in bioreactor

Protein expression was scaled-up in a 1.3 L bioreactor (Eppendorf, USA) with 0.5 L initial volume. Gas flow rate was maintained between 0.5-1.5 vvm (0.5-0.6 Lmin^-1^) by the mass flow controller. The pH of the media was monitored by a pH probe and maintained at 7.0±0.2 by using 3 N phosphoric acid and 12.5% ammonia. Temperature was maintained at 37.0 °C. Dissolved oxygen of the batch was controlled at 30% saturation by cascading the stirrer speed between 300-900 rpm. Bioreactor was monitored and controlled by the Biocommand software (Eppendorf, USA). Fed batch media containing glycerol (200 g L^-1^, v/v) and yeast extract (1%, w/v) was continuously fed to the bacterial culture to enhance the biomass. Protein expression was induced with 1 mM IPTG for 8 h and cells were harvested by centrifugation at 8000 rpm for 15 min. Cell pellet was washed with 0.9% (w/v) NaCl, resuspended in lysis buffer (20 mM Tris-HCL, 150 mM NaCl, 0.5 mM EDTA, pH 8.0), lysed using an Ultrasonicator system (Oscar Ultrsonics Pvt., Ltd., India) for 30 min with 30s on/off (50% duty cycle). Lysed cells were then centrifuged at 7000 rpm for 15 min at 4°C, supernatant was discarded, and the obtained IBs were washed twice with saline. Protein expression was analysed through SDS-PAGE.

### Purification of SARS-CoV-2 N protein

IBs were solubilized in 100 mM Tris-HCl buffer containing 6 M urea (pH 8.0) for 2 hours. The solution was then centrifuged and supernatant was collected. To prepare CEX load, the pH was adjusted to 7.0 using acetic acid and conductivity was adjusted to <3.0 mS/cm using deionised water. The CEX column (SP Sepharose FF, Cytiva USA) was equilibrated with 20 mM phosphate buffer (pH 7.0) and the load was pumped on the CEX column at 5 min retention time. The bound N protein was eluted using 1 M NaCl in 20 mM phosphate buffer (pH 7.0). This purification step also doubled as an on-column refolding step for the N protein. The elute contained a mixture of monomer and dimer of the protein which was separated using preparative SEC (Superdex 200, Cytiva USA). Phosphate buffer (100 mM, pH 7.4) with 10% glycerol (v/v) was used to equilibrate the SEC column and CEX elute (1% of column volume) was injected at 45 min retention time. The SEC output was fractionated, and each fraction was analysed using analytical SEC. Purified monomer and dimer fractions were used for further analysis.

### Immunoblotting

Purified SARS-CoV-2 N protein was electrophoresed on 4-10% SDS-PAGE (Bio-Rad) and stained with Coomassie Brilliant Blue G-250 (CBBG-250). For the detection of N protein through immunoblotting, the obtained protein bands were transferred onto a 0.22 μm nitrocellulose membrane (MDI) using a Trans-Blot Turbo Transfer System (Bio-Rad). The membrane was blocked with 5% (w/v) skimmed milk in Tris buffered saline-0.05% (v/v) Tween-20 (TBST) under gentle shaking at room temperature for 1 h. The membrane was then washed thrice with 1X-TBST and incubated with anti-SARS-CoV-2 N protein antibodies (catalogue No. ab272852, Abcam, 1:1000) in TBST with BSA (2%, w/v). The membrane was then washed thrice with 1X-TBST. Immunoblot was then incubated with HRP conjugated-goat anti-human IgG secondary antibodies (Millipore, AP309P) in a dilution of 1:10,000 for 1 h. The immunoblot was washed thrice with 1X TBST and was visualized on SuperSignal™ West Pico PLUS Chemiluminescent Substrate (Thermo Fischer Scientific), and chemiluminescent signals were captured using ImageQuant LAS 500 instrument (GE Healthcare).

### Peptide mass fingerprinting

In-gel digestion with trypsin protocol was followed for mapping of purified recombinant SARS-CoV-2 N protein^33^. Gel band of interest was excised and transferred into microcentrifuge tube and destained by incubating for 30 minutes in 100 μL of 50 mM ammonium bicarbonate/acetonitrile (1:1, v/v) with vertexing. Then the gel pieces were incubated and vortexed in 200 μL of acetonitrile. Trypsin (Agilent Technologies, California, USA) was added and incubated at 37 °C for 12-14 hrs. Peptides were extracted by adding 100 μL of 1:2 solution of 5% formic acid and acetonitrile and incubated for 15 mins in a shaker at 37 °C. The liquid obtained was evaporated by Speed-Vac vacuum centrifuge and was reconstituted in the 0.1% formic acid for the LC-MS. Digested peptides were separated on a C18 column (Advance Bio Peptide mapping Plus C18, 2.7 μm, 2.1 × 150 mm) using Agilent 1260 HPLC with detector at 214 nm. Column temperature was maintained at 55 °C. The column was equilibrated with 98% solvent A (0.1% TFA in water) and 2% solvent B (0.1% TFA in acetonitrile) for 10 min with a flow rate of 0.5 mlmin^-1^. Elution was achieved with a linear gradient of 2–45% B for 45 min followed by 45–60% B for 10 min, then linear gradient to 100% B for 10 min. Column was cleaned with 100% B for 10 min followed by equilibration with 98% A for 10 min. LC was coupled with ESI-TOF (Agilent Technologies, California USA) and TIC were recorded for m/z 100-3200. The capillary was set at a temperature of 300 °C with a gas flow rate of 8 L/min and nebulizer at 35 psig in positive ion mode. MS spectrum was analysed with Agilent MassHunter Qualitative analysis software (B.07.00).

### Molecular mass identification of monomer and dimer form of N protein

SEC was performed on Dionex Ultimate 3000 HPLC (Thermo Scientific, Sunnyvale, CA, USA) with Superdex 200 (Cytiva, Marlborough, USA) maintained at 25°C. The column was equilibrated with 50 mM phosphate buffer of pH 6.8, 300 mM NaCl salt concentration and 0.02% sodium azide at a flow rate of 0.5 ml/min. Both the fractions of the purified N protein were injected and run for 50 min at a flow rate of 0.5 mL/min and detected at 280 nm. SEC was coupled with MALS from Wyatt technologies, CA, USA, to confirm the molecular mass of the monomer and dimer fractions. All the buffers were filtered through a 0.22 μm membrane (Pall Life Sciences, NY, USA).

### Circular dichroism spectroscopy

Circular dichroism spectra were recorded with a Jasco J-1500 spectrophotometer (Jasco Inc., Maryland, U.S). Secondary structure was measured in the Far-UV range from 195-250 nm with a 1 nm step size. Data were normalized by subtracting the baseline with the buffer and smoothed with Savitzky–Golay smoothing filter.

### Fluorescence spectroscopy

Fluorescence spectroscopy was performed on a Cary Eclipse Fluorescence Spectrophotometer (Agilent Technologies, Santa Clara, California, United States) using Costar 96-well black polystyrene plate. The tryptophan fluorescence was recorded with excitation at 285 nm and emission between 300-500 nm. Slits for both excitation and emission were 5 nm.

### Enzyme linked immunosorbent assay

The fraction of dimeric form of purified N protein of SARS-CoV-2 was diluted at 10 ngµL^-1^ in 0.05 M carbonate-bicarbonate buffer, pH 9.6. The diluted protein was coated in an increasing gradient (25-200 ng well^-1^) on a 96-microtiter ELISA plate (Nunc, Thermo Fisher Scientific) overnight at 4°C. On the subsequent day, unbound protein was removed, and wells were washed thrice with 1X TBST buffer. Wells were then blocked with 4% (w/v) skimmed milk prepared in 1X TBST buffer and incubated at 37°C for 1 h. The anti-SARS-CoV-2 N protein antibodies (Abcam) were diluted in 1X TBST and 100 μL of the diluted antibodies were allowed to interact with the coated N protein in the ELISA wells at 37°C for 1 h. Wells were then washed with 200 μL of 1X TBST buffer three times followed by incubation with 100 μL of goat IgG-HRP antibody (Thermo Fisher Scientific) prepared in 1X TBST buffer. The wells were then washed three times with 200 μL of 1X TBST buffer. One hundred microlitre 3,3′,5,5′-tetramethylbenzidine substrate (Thermo Fisher Scientific) was added to each well and incubated for 10-15 min. The reaction was stopped by adding 100 μL of 0.18 M sulphuric acid and the optical densities of the plate wells were measured using Biotek plate reader at 450 nm.

### Monomeric/dimeric structure modelling

The sequence of SARS-CoV-2 N protein was collected from Uniprot^33^ database with uniport ID: PODTC9. Multi template approach was used to build the model of SARS-CoV-2 N protein. Modeling of the structure was performed using the online version of the HHpred tool^34^. This identified the most promising template for building the structure of the SARS-CoV-2 N protein. Final template-based modeling was performed using the modeller tool^35^. This resulted in monomeric structure of SARS-CoV-2 N protein sequence. Dimeric structure was built using the PDB template 6WZO structure^36^. Pymol tool was used to superimpose the structure of 6WZO and modelled monomeric structure to build its dimeric form.

### Solvent accessible surface area (SASA) calculation

SASA was calculated for monomeric and dimeric form using the naccess tool (http://www.bioinf.manchester.ac.uk/naccess/nacdownload.html).

### Epitopic prediction

Discontinuous epitopes were predicted using the 3D structure of a protein. Monomer and dimer were compared using several tools to identify these discontinuous fragments of the protein that can act as epitopes for antibody binding. Ellipro^37^ was first used for this prediction. The starting residues of 1-48 in the monomer and dimer model protein structure did not appear as globular and were present at the terminal in extended conformation. They were not included in epitope prediction to avoid false positives. Later, a similar analysis was performed with the DiscoTope server. This method predicted the probability of each residue to be part of an epitope.

## Conclusion

A plausible approach to combat pandemic caused by SARS-CoV-2 is to improve the sensitivity of the existing diagnostics. N protein of the SARS-CoV-2 is an important candidate for the development of various diagnosis assays of the COVID-19. This work presents a reliable and sensitive ELISA based diagnostic assay for the detection of antibodies against SARS-CoV-2 N protein. Full length N protein expressed in *E. coli* mainly consist of monomeric and dimeric conformation in the solution. N protein monomer and dimer conformations were characterized by CD suggesting improved secondary structure of the dimer due to oligomerization. Similarly, fluorescence spectroscopy indicates the exposure of buried tryptophan in dimer resulting in the oligomerization. Indirect ELISA developed by using purified monomer and dimer N protein showed enhanced sensitivity of the N protein dimer as compared to the monomer. We further confirm this observation by SASA, which predict stabilization of N protein dimer by the burial of hydrophobic residues. Our findings have significantly upgraded the present understanding of SAR-CoV-2 N protein and its application in diagnostic assays. We further believe that the employment of N protein dimer in the diagnostic assays will improve the sensitivity of the existing assays.

## Acknowledgments

This work was funded by a Center of Excellence for Biopharmaceutical Technology grant from the Department of Biotechnology, Government of India (BT/COE/34/SP15097/2015) and CSR funding from Microsoft Corporation.

## Author contributions

A.S.R., W.H.K., R.B. designed the study. W.H.K., N.K., S.G., V.B., and D.M performed the experiments. W.H.K., N.K., S.G, V.B., A.M., D.M. and R.B. analysed and interpreted data. A.M. performed the bioinformatics work. W.H.K., N.K., S.G., V.B., and A.M., wrote the original manuscript. A.S.R. supervised the project and was the recipient of the funding. All authors were involved in the drafting, review, and approval of the report and the decision to submit for publication.

## Additional information

Supplementary information of Table S1 is attached with the manuscript.

## References

1. M. Iyer, K. Jayaramayya, M. D. Subramaniam, S. B. Lee, A. A. Dayem, S. G. Cho and B. Vellingiri, BMB Rep, 2020, 53, 191–205.

2. V. Kumar, K. U. Doshi, W. H. Khan and A. S. Rathore, Journal of Chemical Technology & Biotechnology, 2021, 96, 299–308.

3. M. Zeyaullah, A. M Alshahrani, K. Muzammil, I. Ahmad, S. Alam, W. H. Khan, R. Ahmad, Frontiers in Genetics, 2021, DOI: 10.3389/fgene.2021.693916.

4. J. B. Mahony, Clinical Microbiology Reviews, 2008, 21, 716–747.

5. Z. R. Tehrani, S. Saadat, E. Saleh, X. Ouyang, N. Constantine, A. L. DeVico, A. D. Harris, G. K. Lewis, S. Kottilil and M. M. Sajadi, medRxiv, 2020, DOI: 10.1101/2020.08.05.20168476, 2020.2008.2005.20168476.

6. R. Yuen, D. Steiner, R. Pihl, E. Chavez, A. Olson, L. Baird, F. Korkmaz, P. Urick, M. Sagar, J. Berrigan, R. Gummuluru, R. Corley, K. Quillen, A. Belkina, G. Mostoslavsky, I. Rifkin, Y. Kataria, A. Cappione, N. Lin, N. Bhadelia and J. Snyder-Cappione, medRxiv, 2020, DOI: 10.1101/2020.09.15.20192765, 2020.2009.2015.20192765.

7. C. K. Chang, M. H. Hou, C. F. Chang, C. D. Hsiao and T. H. Huang, Antiviral Res, 2014, 103, 39–50.

8. S. Satarker and M. Nampoothiri, Archives of medical research, 2020, 51, 482–491.

9. C. Wu, A. J. Qavi, A. Hachim, N. Kavian, A. R. Cole, A. B. Moyle, N. D. Wagner, J. Sweeney-Gibbons, H. W. Rohrs, M. L. Gross, J. S. M. Peiris, C. F. Basler, C. W. Farnsworth, S. A. Valkenburg, G. K. Amarasinghe and D. W. Leung, bioRxiv, 2020, DOI: 10.1101/2020.11.30.404905, 2020.2011.2030.404905.

10. C. Y. Chen, C. K. Chang, Y. W. Chang, S. C. Sue, H. I. Bai, L. Riang, C. D. Hsiao and T. H. Huang, J Mol Biol, 2007, 368, 1075–1086.

11. H. Jayaram, H. Fan, B. R. Bowman, A. Ooi, J. Jayaram, E. W. Collisson, J. Lescar and B. V. V. Prasad, Journal of Virology, 2006, 80, 6612–6620.

12. M. Takeda, C. K. Chang, T. Ikeya, P. Güntert, Y. H. Chang, Y. L. Hsu, T. H. Huang and M. Kainosho, J Mol Biol, 2008, 380, 608–622.

13. I. M. Yu, M. L. Oldham, J. Zhang and J. Chen, J Biol Chem, 2006, 281, 17134–17139.

14. T. H. V. Nguyen, J. Lichière, B. Canard, N. Papageorgiou, S. Attoumani, F. Ferron and B. Coutard, Acta crystallographica. Section D, Structural biology, 2019, 75, 8–15.

15. H. Luo, J. Chen, K. Chen, X. Shen and H. Jiang, Biochemistry, 2006, 45, 11827–11835.

16. G. O. Sabbih, M. A. Korsah, J. Jeevanandam and M. K. Danquah, Biotechnol Prog, 2020, DOI: 10.1002/btpr.3096e3096BTPR3096 [pii], e3096.

17. Q. Ye, A. M. V. West, S. Silletti and K. D. Corbett, bioRxiv, 2020, DOI: 10.1101/2020.05.17.100685, 2020.2005.2017.100685.

18. L. F. Huergo, M. S. Conzentino, E. C. Gerhardt, A. R. Santos, F. de Oliveira Pedrosa, E. M. Souza, M. B. Nogueira, K. Forchhammer, F. G. Rego and S. M. Raboni, medRxiv, 2020.

19. K. A. Timani, L. Ye, L. Ye, Y. Zhu, Z. Wu and Z. Gong, Journal of clinical virology, 2004, 30, 309–312.

20. N. K. Dutta, K. Mazumdar, B. H. Lee, M. W. Baek, D. J. Kim, Y. R. Na, S. H. Park, H. K. Lee, H. Kariwa, Q. Mai le and J. H. Park, Immunol Lett, 2008, 118, 65–71.

21. F. Bonelli, A. Sarasini, C. Zierold, M. Calleri, A. Bonetti, C. Vismara, F. A. Blocki, L. Pallavicini, A. Chinali, D. Campisi, E. Percivalle, A. P. DiNapoli, C. F. Perno and F. Baldanti, Journal of Clinical Microbiology, 2020, 58, e01224–01220.

22. B. Diego-Martin, B. González, M. Vazquez-Vilar, S. Selma, R. Mateos-Fernández, S. Gianoglio, A. Fernández-del-Carmen and D. Orzáez, bioRxiv, 2020, DOI: 10.1101/2020.10.13.331306, 2020.2010.2013.331306.

23. H. K. Lee, B. H. Lee, N. K. Dutta, S. H. Seok, M. W. Baek, H. Y. Lee, D. J. Kim, Y. R. Na, K. J. Noh, S. H. Park, H. Kariwa, M. Nakauchi, Q. Mai le, S. J. Heo and J. H. Park, J Microbiol Biotechnol, 2008, 18, 1717–1721.

24. W. Liu, L. Liu, G. Kou, Y. Zheng, Y. Ding, W. Ni, Q. Wang, L. Tan, W. Wu, S. Tang, Z. Xiong and S. Zheng, Journal of Clinical Microbiology, 2020, 58, e00461–00420.

25. M. S. Makatsa, M. B. Tincho, J. M. Wendoh, S. D. Ismail, R. Nesamari, F. Pera, S. de Beer, A. David, S. Jugwanth, M. P. Gededzha, N. Mampeule, I. Sanne, W. Stevens, L. Scott, J. Blackburn, E. S. Mayne, R. S. Keeton and W. A. Burgers, medRxiv, 2020, DOI: 10.1101/2020.08.04.20167940, 2020.2008.2004.20167940.

26. L. Mazzini, D. Martinuzzi, I. Hyseni, G. Lapini, L. Benincasa, P. Piu, C. M. Trombetta, S. Marchi, I. Razzano, A. Manenti and E. Montomoli, bioRxiv, 2020, DOI: 10.1101/2020.08.10.243717, 2020.2008.2010.243717.

27. R. Yokoyama, M. Kurano, Y. Morita, T. Shimura, Y. Nakano, C. Qian, F. Xia, F. He, Y. Kishi, J. Okada, N. Yoshikawa, Y. Nagura, H. Okazaki, K. Moriya, Y. Seto, T. Kodama and Y. Yatomi, medRxiv, 2020, DOI: 10.1101/2020.07.16.20155796, 2020.2007.2016.20155796.

28. L. Liu, W. Liu, Y. Zheng, X. Jiang, G. Kou, J. Ding, Q. Wang, Q. Huang, Y. Ding, W. Ni, W. Wu, S. Tang, L. Tan, Z. Hu, W. Xu, Y. Zhang, B. Zhang, Z. Tang, X. Zhang, H. Li, Z. Rao, H. Jiang, X. Ren, S. Wang and S. Zheng, Microbes Infect, 2020, 22, 206–211.

29. H. E. Prince, T. S. Givens, M. Lapé-Nixon, N. J. Clarke, D. A. Schwab, H. J. Batterman, R. S. Jones, W. A. Meyer, H. Kapoor, C. M. Rowland, F. Haji-Sheikhi and E. M. Marlowe, Journal of Clinical Microbiology, 2020, 58, e01742–01720.

30. W. Zeng, G. Liu, H. Ma, D. Zhao, Y. Yang, M. Liu, A. Mohammed, C. Zhao, Y. Yang and J. Xie, Biochemical and biophysical research communications, 2020, 527, 618–623.

31. P. D. Burbelo, F. X. Riedo, C. Morishima, S. Rawlings, D. Smith, S. Das, J. R. Strich, D. S. Chertow, R. T. Davey, Jr. and J. I. Cohen, medRxiv, 2020, DOI: 2020.04.20.20071423 [pii]10.1101/2020.04.20.20071423.

32. C. A. Lutomski, T. J. El-Baba, J. R. Bolla and C. V. Robinson, bioRxiv, 2020, DOI: 10.1101/2020.10.06.328112, 2020.2010.2006.328112.33..

34. J. Söding, A. Biegert and A. N. Lupas, Nucleic acids research, 2005, 33, W244–W248.

35. B. Webb and A. Sali, Current protocols in bioinformatics, 2016, 54, 5.6. 1-5.6. 37.

36. Q. Ye, A. M. West, S. Silletti and K. D. Corbett, Protein Science, 2020, 29, 1890–1901.

37. J. Ponomarenko, H.-H. Bui, W. Li, N. Fusseder, P. E. Bourne, A. Sette and B. Peters, BMC bioinformatics, 2008, 9, 1–8.

