## Supplemental table 1 for "Dimerization of SARS-CoV-2 nucleocapsid protein affects sensitivity of ELISA based diagnostics of COVID-19"

**Supplementary Table 1:** Residue epitope propensity score for monomer and dimer calculated using DiscoTope server. Only epitopic residues (<=B) are listed in the table.

**Monomer**

| Residue # | Residue Name | Contact Number | Propensity Score | DiscoTope Score | Identified B cell epitope |
| --- | --- | --- | --- | --- | --- |
| 9 | GLN | 2 | 2.932 | 2.365 | <=B |
| 10 | ARG | 3 | 2.733 | 2.073 | <=B |
| 11 | ASN | 3 | 2.569 | 1.928 | <=B |
| 12 | ALA | 3 | 2.048 | 1.467 | <=B |
| 13 | PRO | 0 | 2.326 | 2.058 | <=B |
| 14 | ARG | 0 | 1.976 | 1.749 | <=B |
| 15 | ILE | 6 | 1.679 | 0.796 | <=B |
| 16 | THR | 3 | 1.282 | 0.79 | <=B |
| 17 | PHE | 1 | 0.802 | 0.595 | <=B |
| 18 | GLY | 2 | 0.683 | 0.374 | <=B |
| 19 | GLY | 5 | 0.925 | 0.244 | <=B |
| 20 | PRO | 3 | 1.421 | 0.913 | <=B |
| 21 | SER | 3 | 1.195 | 0.712 | <=B |
| 22 | ASP | 2 | 1.533 | 1.127 | <=B |
| 23 | SER | 2 | 2.351 | 1.85 | <=B |
| 24 | THR | 1 | 2.517 | 2.113 | <=B |
| 25 | GLY | 5 | 3.05 | 2.124 | <=B |
| 26 | SER | 2 | 3.693 | 3.039 | <=B |
| 27 | ASN | 4 | 4.133 | 3.197 | <=B |
| 28 | GLN | 0 | 4.185 | 3.704 | <=B |
| 29 | ASN | 3 | 3.762 | 2.984 | <=B |
| 30 | GLY | 6 | 3.673 | 2.56 | <=B |
| 31 | GLU | 3 | 3.38 | 2.646 | <=B |
| 32 | ARG | 0 | 3.283 | 2.905 | <=B |
| 33 | SER | 4 | 2.989 | 2.185 | <=B |
| 34 | GLY | 3 | 2.736 | 2.076 | <=B |
| 35 | ALA | 1 | 2.697 | 2.272 | <=B |
| 36 | ARG | 3 | 3.429 | 2.69 | <=B |
| 37 | SER | 3 | 4.194 | 3.367 | <=B |
| 38 | LYS | 3 | 4.959 | 4.043 | <=B |
| 39 | GLN | 3 | 5.674 | 4.677 | <=B |
| 40 | ARG | 3 | 6.367 | 5.29 | <=B |
| 41 | ARG | 3 | 7.114 | 5.951 | <=B |
| 42 | PRO | 3 | 6.961 | 5.815 | <=B |
| 43 | GLN | 3 | 6.445 | 5.359 | <=B |
| 44 | GLY | 0 | 5.644 | 4.995 | <=B |
| 45 | LEU | 8 | 4.2 | 2.797 | <=B |
| 46 | PRO | 0 | 4.388 | 3.883 | <=B |
| 47 | ASN | 15 | 3.99 | 1.806 | <=B |
| 48 | ASN | 8 | 1.43 | 0.346 | <=B |
| 93 | ARG | 8 | 3.503 | 2.18 | <=B |
| 94 | ILE | 6 | 2.842 | 1.825 | <=B |
| 95 | ARG | 21 | 3.583 | 0.756 | <=B |
| 96 | GLY | 1 | 5.224 | 4.508 | <=B |
| 97 | GLY | 20 | 4.124 | 1.35 | <=B |
| 98 | ASP | 2 | 3.303 | 2.693 | <=B |
| 101 | MET | 15 | 3.938 | 1.76 | <=B |
| 102 | LYS | 15 | 3.055 | 0.979 | <=B |
| 103 | ASP | 18 | 2.456 | 0.103 | <=B |
| 120 | GLY | 3 | 1.862 | 1.303 | <=B |
| 122 | PRO | 2 | 0.44 | 0.159 | <=B |
| 143 | LYS | 10 | 2.553 | 1.109 | <=B |
| 145 | HIS | 10 | 1.721 | 0.373 | <=B |
| 183 | ALA | 9 | 1.174 | 0.004 | <=B |
| 189 | LYS | 20 | 2.693 | 0.084 | <=B |
| 191 | ARG | 21 | 3.497 | 0.68 | <=B |
| 192 | GLN | 17 | 4.375 | 1.917 | <=B |
| 193 | LYS | 8 | 3.302 | 2.002 | <=B |
| 194 | ARG | 25 | 3.576 | 0.29 | <=B |
| 195 | THR | 4 | 3.049 | 2.239 | <=B |
| 196 | ALA | 21 | 3.251 | 0.462 | <=B |
| 197 | THR | 22 | 3.302 | 0.392 | <=B |
| 200 | TYR | 14 | 2.989 | 1.036 | <=B |
| 201 | ASN | 10 | 5.112 | 3.374 | <=B |
| 202 | VAL | 25 | 3.702 | 0.401 | <=B |
| 203 | THR | 5 | 5.182 | 4.011 | <=B |
| 204 | GLN | 19 | 5.213 | 2.428 | <=B |
| 205 | ALA | 26 | 4.603 | 1.084 | <=B |
| 206 | PHE | 28 | 4.591 | 0.843 | <=B |
| 207 | GLY | 8 | 4.468 | 3.034 | <=B |
| 208 | ARG | 8 | 4.62 | 3.169 | <=B |
| 209 | ARG | 25 | 5.26 | 1.78 | <=B |
| 210 | GLY | 8 | 3.652 | 2.312 | <=B |
| 211 | PRO | 3 | 3.189 | 2.478 | <=B |
| 212 | GLU | 6 | 5.031 | 3.762 | <=B |
| 213 | GLN | 6 | 4.444 | 3.243 | <=B |
| 214 | THR | 9 | 4.087 | 2.582 | <=B |
| 215 | GLN | 18 | 3.445 | 0.979 | <=B |
| 217 | ASN | 11 | 3.849 | 2.142 | <=B |
| 220 | ASP | 14 | 3.95 | 1.885 | <=B |
| 221 | GLN | 4 | 3.965 | 3.049 | <=B |
| 222 | GLU | 10 | 3.836 | 2.245 | <=B |
| 223 | LEU | 27 | 3.946 | 0.387 | <=B |
| 224 | ILE | 21 | 4.773 | 1.81 | <=B |
| 225 | ARG | 6 | 5.036 | 3.767 | <=B |
| 226 | GLN | 11 | 4.484 | 2.703 | <=B |
| 227 | GLY | 20 | 3.645 | 0.926 | <=B |
| 228 | THR | 9 | 1.946 | 0.688 | <=B |
| 229 | ASP | 14 | 3.497 | 1.485 | <=B |
| 231 | LYS | 5 | 2.495 | 1.633 | <=B |
| 233 | TRP | 18 | 2.418 | 0.07 | <=B |
| 234 | PRO | 2 | 1.329 | 0.946 | <=B |
| 273 | ASP | 0 | 1.459 | 1.291 | <=B |
| 274 | LYS | 2 | 2.161 | 1.683 | <=B |
| 275 | ASP | 13 | 2.585 | 0.793 | <=B |
| 276 | PRO | 0 | 3.295 | 2.916 | <=B |
| 277 | ASN | 13 | 2.313 | 0.552 | <=B |
| 278 | PHE | 8 | 1.246 | 0.183 | <=B |
| 279 | LYS | 1 | 1.774 | 1.455 | <=B |
| 280 | ASP | 11 | 2.066 | 0.563 | <=B |
| 283 | ILE | 6 | 1.471 | 0.612 | <=B |
| 286 | ASN | 6 | 2.272 | 1.321 | <=B |
| 287 | LYS | 16 | 2.996 | 0.811 | <=B |
| 289 | ILE | 7 | 2.408 | 1.326 | <=B |
| 290 | ASP | 6 | 1.99 | 1.071 | <=B |
| 292 | TYR | 16 | 3.699 | 1.433 | <=B |
| 293 | LYS | 6 | 3.797 | 2.671 | <=B |
| 294 | THR | 12 | 3.538 | 1.751 | <=B |
| 295 | PHE | 19 | 3.115 | 0.572 | <=B |
| 296 | PRO | 2 | 3.287 | 2.679 | <=B |

**Dimer**

| Residue # | Residue Name | Contact Number | Propensity Score | DiscoTope Score | Identified B cell epitope |
| --- | --- | --- | --- | --- | --- |
| 9 | GLN | 2 | 2.932 | 2.365 | <=B |
| 10 | ARG | 3 | 2.733 | 2.073 | <=B |
| 11 | ASN | 3 | 2.569 | 1.928 | <=B |
| 12 | ALA | 3 | 2.048 | 1.467 | <=B |
| 13 | PRO | 0 | 2.326 | 2.058 | <=B |
| 14 | ARG | 0 | 1.976 | 1.749 | <=B |
| 15 | ILE | 6 | 1.679 | 0.796 | <=B |
| 16 | THR | 3 | 1.282 | 0.79 | <=B |
| 17 | PHE | 1 | 0.802 | 0.595 | <=B |
| 18 | GLY | 2 | 0.683 | 0.374 | <=B |
| 19 | GLY | 5 | 0.925 | 0.244 | <=B |
| 20 | PRO | 3 | 1.421 | 0.913 | <=B |
| 21 | SER | 3 | 1.195 | 0.712 | <=B |
| 22 | ASP | 2 | 1.533 | 1.127 | <=B |
| 23 | SER | 2 | 2.351 | 1.85 | <=B |
| 24 | THR | 1 | 2.517 | 2.113 | <=B |
| 25 | GLY | 5 | 3.05 | 2.124 | <=B |
| 26 | SER | 2 | 3.693 | 3.039 | <=B |
| 27 | ASN | 4 | 4.133 | 3.197 | <=B |
| 28 | GLN | 0 | 4.185 | 3.704 | <=B |
| 29 | ASN | 3 | 3.762 | 2.984 | <=B |
| 30 | GLY | 6 | 3.673 | 2.56 | <=B |
| 31 | GLU | 3 | 3.38 | 2.646 | <=B |
| 32 | ARG | 0 | 3.283 | 2.905 | <=B |
| 33 | SER | 4 | 2.989 | 2.185 | <=B |
| 34 | GLY | 3 | 2.736 | 2.076 | <=B |
| 35 | ALA | 1 | 2.697 | 2.272 | <=B |
| 36 | ARG | 3 | 3.429 | 2.69 | <=B |
| 37 | SER | 3 | 4.43 | 3.575 | <=B |
| 38 | LYS | 3 | 5.267 | 4.316 | <=B |
| 39 | GLN | 3 | 6.082 | 5.037 | <=B |
| 40 | ARG | 3 | 7.789 | 6.548 | <=B |
| 41 | ARG | 3 | 8.1 | 6.824 | <=B |
| 42 | PRO | 3 | 8.054 | 6.783 | <=B |
| 43 | GLN | 3 | 7.498 | 6.291 | <=B |
| 44 | GLY | 0 | 7.33 | 6.487 | <=B |
| 45 | LEU | 10 | 6.353 | 4.473 | <=B |
| 46 | PRO | 6 | 6.831 | 5.355 | <=B |
| 47 | ASN | 15 | 6.018 | 3.601 | <=B |
| 79 | SER | 15 | 2.852 | 0.799 | <=B |
| 81 | ASP | 6 | 1.719 | 0.832 | <=B |
| 93 | ARG | 8 | 4.229 | 2.823 | <=B |
| 94 | ILE | 13 | 4.452 | 2.445 | <=B |
| 95 | ARG | 29 | 6.211 | 2.162 | <=B |
| 96 | GLY | 3 | 9.796 | 8.324 | <=B |
| 97 | GLY | 20 | 8.815 | 5.501 | <=B |
| 98 | ASP | 3 | 8.534 | 7.208 | <=B |
| 99 | GLY | 19 | 6.056 | 3.175 | <=B |
| 100 | LYS | 18 | 4.783 | 2.163 | <=B |
| 101 | MET | 15 | 5.796 | 3.405 | <=B |
| 102 | LYS | 17 | 4.642 | 2.153 | <=B |
| 103 | ASP | 18 | 3.076 | 0.652 | <=B |
| 120 | GLY | 3 | 1.92 | 1.354 | <=B |
| 143 | LYS | 10 | 3.938 | 2.335 | <=B |
| 145 | HIS | 13 | 4.265 | 2.28 | <=B |
| 147 | GLY | 10 | 3.74 | 2.16 | <=B |
| 183 | ALA | 9 | 1.742 | 0.506 | <=B |
| 185 | GLU | 6 | 0.815 | 0.031 | <=B |
| 188 | LYS | 27 | 3.517 | 0.008 | <=B |
| 189 | LYS | 21 | 4.865 | 1.891 | <=B |
| 190 | PRO | 26 | 3.52 | 0.125 | <=B |
| 191 | ARG | 21 | 4.898 | 1.92 | <=B |
| 192 | GLN | 18 | 7.903 | 4.924 | <=B |
| 193 | LYS | 8 | 6.461 | 4.798 | <=B |
| 194 | ARG | 25 | 5.506 | 1.998 | <=B |
| 195 | THR | 4 | 3.88 | 2.973 | <=B |
| 196 | ALA | 21 | 3.975 | 1.103 | <=B |
| 197 | THR | 22 | 3.534 | 0.598 | <=B |
| 200 | TYR | 14 | 3.841 | 1.789 | <=B |
| 201 | ASN | 10 | 6.47 | 4.576 | <=B |
| 202 | VAL | 25 | 4.947 | 1.503 | <=B |
| 203 | THR | 5 | 7.033 | 5.649 | <=B |
| 204 | GLN | 19 | 7.331 | 4.303 | <=B |
| 205 | ALA | 26 | 7.035 | 3.236 | <=B |
| 206 | PHE | 28 | 6.851 | 2.843 | <=B |
| 207 | GLY | 12 | 6.787 | 4.627 | <=B |
| 208 | ARG | 9 | 6.335 | 4.571 | <=B |
| 209 | ARG | 25 | 6.463 | 2.845 | <=B |
| 210 | GLY | 8 | 4.631 | 3.179 | <=B |
| 211 | PRO | 3 | 3.617 | 2.856 | <=B |
| 212 | GLU | 6 | 6.291 | 4.878 | <=B |
| 213 | GLN | 6 | 5.161 | 3.877 | <=B |
| 214 | THR | 11 | 6.368 | 4.371 | <=B |
| 215 | GLN | 18 | 5.302 | 2.623 | <=B |
| 216 | GLY | 23 | 3.54 | 0.488 | <=B |
| 217 | ASN | 11 | 4.05 | 2.32 | <=B |
| 220 | ASP | 14 | 3.95 | 1.885 | <=B |
| 221 | GLN | 4 | 3.965 | 3.049 | <=B |
| 222 | GLU | 10 | 3.836 | 2.245 | <=B |
| 223 | LEU | 27 | 3.946 | 0.387 | <=B |
| 224 | ILE | 21 | 4.995 | 2.006 | <=B |
| 225 | ARG | 6 | 5.002 | 3.736 | <=B |
| 226 | GLN | 11 | 4.484 | 2.703 | <=B |
| 227 | GLY | 20 | 3.645 | 0.926 | <=B |
| 228 | THR | 9 | 1.946 | 0.688 | <=B |
| 229 | ASP | 14 | 3.497 | 1.485 | <=B |
| 231 | LYS | 5 | 2.495 | 1.633 | <=B |
| 233 | TRP | 18 | 2.418 | 0.07 | <=B |
| 234 | PRO | 2 | 1.329 | 0.946 | <=B |
| 253 | GLY | 8 | 1.224 | 0.163 | <=B |
| 262 | TRP | 13 | 1.908 | 0.193 | <=B |
| 273 | ASP | 0 | 1.459 | 1.291 | <=B |
| 274 | LYS | 2 | 2.161 | 1.683 | <=B |
| 275 | ASP | 13 | 2.585 | 0.793 | <=B |
| 276 | PRO | 0 | 3.295 | 2.916 | <=B |
| 277 | ASN | 13 | 2.313 | 0.552 | <=B |
| 278 | PHE | 8 | 1.246 | 0.183 | <=B |
| 279 | LYS | 1 | 1.774 | 1.455 | <=B |
| 280 | ASP | 11 | 2.066 | 0.563 | <=B |
| 283 | ILE | 6 | 1.471 | 0.612 | <=B |
| 286 | ASN | 6 | 2.272 | 1.321 | <=B |
| 287 | LYS | 16 | 2.996 | 0.811 | <=B |
| 289 | ILE | 7 | 2.408 | 1.326 | <=B |
| 290 | ASP | 6 | 1.99 | 1.071 | <=B |
| 292 | TYR | 16 | 3.699 | 1.433 | <=B |
| 293 | LYS | 6 | 3.797 | 2.671 | <=B |
| 294 | THR | 12 | 3.538 | 1.751 | <=B |
| 295 | PHE | 19 | 3.115 | 0.572 | <=B |
| 296 | PRO | 2 | 3.287 | 2.679 | <=B |
| 9 | GLN | 2 | 2.932 | 2.365 | <=B |
| 10 | ARG | 3 | 2.733 | 2.073 | <=B |
| 11 | ASN | 3 | 2.569 | 1.928 | <=B |
| 12 | ALA | 3 | 2.048 | 1.467 | <=B |
| 13 | PRO | 0 | 2.326 | 2.058 | <=B |
| 14 | ARG | 0 | 1.976 | 1.749 | <=B |
| 15 | ILE | 6 | 1.679 | 0.796 | <=B |
| 16 | THR | 3 | 1.282 | 0.79 | <=B |
| 17 | PHE | 1 | 0.802 | 0.595 | <=B |
| 18 | GLY | 2 | 0.683 | 0.374 | <=B |
| 19 | GLY | 5 | 0.925 | 0.244 | <=B |
| 20 | PRO | 3 | 1.421 | 0.913 | <=B |
| 21 | SER | 3 | 1.195 | 0.712 | <=B |
| 22 | ASP | 2 | 1.533 | 1.126 | <=B |
| 23 | SER | 2 | 2.351 | 1.85 | <=B |
| 24 | THR | 1 | 2.517 | 2.113 | <=B |
| 25 | GLY | 5 | 3.05 | 2.124 | <=B |
| 26 | SER | 2 | 3.693 | 3.039 | <=B |
| 27 | ASN | 4 | 4.133 | 3.197 | <=B |
| 28 | GLN | 0 | 4.185 | 3.704 | <=B |
| 29 | ASN | 3 | 3.762 | 2.984 | <=B |
| 30 | GLY | 6 | 3.673 | 2.56 | <=B |
| 31 | GLU | 3 | 3.38 | 2.646 | <=B |
| 32 | ARG | 0 | 3.283 | 2.905 | <=B |
| 33 | SER | 4 | 2.989 | 2.185 | <=B |
| 34 | GLY | 3 | 2.736 | 2.076 | <=B |
| 35 | ALA | 1 | 2.697 | 2.272 | <=B |
| 36 | ARG | 3 | 3.429 | 2.69 | <=B |
| 37 | SER | 3 | 4.43 | 3.575 | <=B |
| 38 | LYS | 3 | 5.267 | 4.316 | <=B |
| 39 | GLN | 3 | 6.081 | 5.037 | <=B |
| 40 | ARG | 3 | 7.789 | 6.548 | <=B |
| 41 | ARG | 3 | 8.1 | 6.824 | <=B |
| 42 | PRO | 3 | 8.054 | 6.782 | <=B |
| 43 | GLN | 3 | 7.498 | 6.291 | <=B |
| 44 | GLY | 0 | 7.33 | 6.487 | <=B |
| 45 | LEU | 10 | 6.353 | 4.473 | <=B |
| 46 | PRO | 6 | 6.831 | 5.355 | <=B |
| 47 | ASN | 15 | 6.018 | 3.601 | <=B |
| 79 | SER | 15 | 2.852 | 0.799 | <=B |
| 81 | ASP | 6 | 1.719 | 0.832 | <=B |
| 93 | ARG | 8 | 4.229 | 2.823 | <=B |
| 94 | ILE | 13 | 4.452 | 2.445 | <=B |
| 95 | ARG | 29 | 6.211 | 2.162 | <=B |
| 96 | GLY | 3 | 9.796 | 8.324 | <=B |
| 97 | GLY | 20 | 8.815 | 5.502 | <=B |
| 98 | ASP | 3 | 8.534 | 7.208 | <=B |
| 99 | GLY | 19 | 6.056 | 3.175 | <=B |
| 100 | LYS | 18 | 4.783 | 2.163 | <=B |
| 101 | MET | 15 | 5.796 | 3.405 | <=B |
| 102 | LYS | 17 | 4.642 | 2.154 | <=B |
| 103 | ASP | 18 | 3.076 | 0.652 | <=B |
| 120 | GLY | 3 | 1.687 | 1.148 | <=B |
| 143 | LYS | 10 | 3.938 | 2.335 | <=B |
| 145 | HIS | 13 | 4.265 | 2.279 | <=B |
| 147 | GLY | 11 | 3.741 | 2.046 | <=B |
| 183 | ALA | 9 | 1.742 | 0.506 | <=B |
| 185 | GLU | 6 | 0.815 | 0.032 | <=B |
| 188 | LYS | 27 | 3.517 | 0.008 | <=B |
| 189 | LYS | 21 | 4.865 | 1.891 | <=B |
| 190 | PRO | 26 | 3.52 | 0.125 | <=B |
| 191 | ARG | 21 | 4.898 | 1.92 | <=B |
| 192 | GLN | 18 | 7.903 | 4.924 | <=B |
| 193 | LYS | 8 | 6.461 | 4.798 | <=B |
| 194 | ARG | 25 | 5.506 | 1.998 | <=B |
| 195 | THR | 4 | 3.88 | 2.973 | <=B |
| 196 | ALA | 21 | 3.975 | 1.103 | <=B |
| 197 | THR | 22 | 3.534 | 0.597 | <=B |
| 200 | TYR | 14 | 3.841 | 1.789 | <=B |
| 201 | ASN | 10 | 6.47 | 4.576 | <=B |
| 202 | VAL | 25 | 4.947 | 1.504 | <=B |
| 203 | THR | 5 | 7.033 | 5.649 | <=B |
| 204 | GLN | 19 | 7.331 | 4.303 | <=B |
| 205 | ALA | 26 | 7.035 | 3.236 | <=B |
| 206 | PHE | 28 | 6.851 | 2.843 | <=B |
| 207 | GLY | 12 | 6.787 | 4.627 | <=B |
| 208 | ARG | 9 | 6.334 | 4.571 | <=B |
| 209 | ARG | 25 | 6.463 | 2.845 | <=B |
| 210 | GLY | 8 | 4.631 | 3.178 | <=B |
| 211 | PRO | 3 | 3.617 | 2.856 | <=B |
| 212 | GLU | 6 | 6.291 | 4.877 | <=B |
| 213 | GLN | 6 | 5.16 | 3.877 | <=B |
| 214 | THR | 11 | 6.368 | 4.371 | <=B |
| 215 | GLN | 18 | 5.302 | 2.623 | <=B |
| 216 | GLY | 23 | 3.54 | 0.488 | <=B |
| 217 | ASN | 11 | 4.05 | 2.319 | <=B |
| 220 | ASP | 14 | 3.949 | 1.885 | <=B |
| 221 | GLN | 4 | 3.965 | 3.049 | <=B |
| 222 | GLU | 10 | 3.836 | 2.245 | <=B |
| 223 | LEU | 27 | 3.946 | 0.387 | <=B |
| 224 | ILE | 21 | 4.995 | 2.006 | <=B |
| 225 | ARG | 6 | 5.002 | 3.736 | <=B |
| 226 | GLN | 11 | 4.484 | 2.703 | <=B |
| 227 | GLY | 20 | 3.645 | 0.926 | <=B |
| 228 | THR | 9 | 1.946 | 0.687 | <=B |
| 229 | ASP | 14 | 3.497 | 1.485 | <=B |
| 231 | LYS | 5 | 2.495 | 1.633 | <=B |
| 233 | TRP | 18 | 2.418 | 0.07 | <=B |
| 234 | PRO | 2 | 1.328 | 0.946 | <=B |
| 253 | GLY | 8 | 1.224 | 0.163 | <=B |
| 262 | TRP | 13 | 1.908 | 0.193 | <=B |
| 273 | ASP | 0 | 1.459 | 1.291 | <=B |
| 274 | LYS | 2 | 2.161 | 1.683 | <=B |
| 275 | ASP | 13 | 2.586 | 0.793 | <=B |
| 276 | PRO | 0 | 3.295 | 2.916 | <=B |
| 277 | ASN | 13 | 2.313 | 0.552 | <=B |
| 278 | PHE | 8 | 1.246 | 0.183 | <=B |
| 279 | LYS | 1 | 1.774 | 1.455 | <=B |
| 280 | ASP | 11 | 2.066 | 0.563 | <=B |
| 283 | ILE | 6 | 1.471 | 0.612 | <=B |
| 286 | ASN | 6 | 2.272 | 1.321 | <=B |
| 287 | LYS | 16 | 2.996 | 0.811 | <=B |
| 289 | ILE | 7 | 2.408 | 1.326 | <=B |
| 290 | ASP | 6 | 1.99 | 1.071 | <=B |
| 292 | TYR | 16 | 3.698 | 1.433 | <=B |
| 293 | LYS | 6 | 3.797 | 2.67 | <=B |
| 294 | THR | 12 | 3.538 | 1.751 | <=B |
| 295 | PHE | 19 | 3.115 | 0.572 | <=B |
| 296 | PRO | 2 | 3.287 | 2.679 | <=B |
